# Host sex and genotype modify the gut microbiome response to helminth infection

**DOI:** 10.1101/608638

**Authors:** Fei Ling, Natalie Steinel, Jesse Weber, Lei Ma, Chris Smith, Decio Correa, Bin Zhu, Daniel Bolnick, Gaoxue Wang

## Abstract

The microbial community can be altered by direct/indirect interactions with parasites infecting host. Direct interactions can arise from physical/chemical contact with the parasite. Indirect interactions can involve parasite-induced changes in host immunity. If so, this would represent a case of genetic polymorphism in one species controlling an ecological interaction between other species. Here, we report a test of this expectation: we experimentally exposed *Gasterosteus aculeatus* to their naturally co-evolved parasite, *Schistocephalus solidus*. The host microbiome differed in response to parasite exposure, and between infected and uninfected fish. The microbial response to infection differed between host sexes, and also varied between variants at autosomal quantitative trait loci (QTL). These results indicate that host genotype regulates the indirect effect of infection on a vertebrate gut microbiome. Our results also raise the possibility that this sex-bias may be related to sex-specific microbial responses to the presence (or, absence) of helminthes. Therefore, helminth-based therapeutics as possible treatments for inflammatory bowel diseases might need to take account of these interactions, potentially requiring therapies tailored to host sex or genotype.

## Introduction

Helminth infection is often associated with changes in the host’s gut microbiome [1, 2], resulting in a complex multi-way ecological interaction between host, parasite, and diverse microbiota. For example, *Hymenolepis diminuta* infection reduces the abundance of Bacilli species in the gut of mammalian hosts, in both the lab and wild [3, 4]. However, other studies report no effects of infection on the microbiome: *Trichuris trichiura* and *Necator americanus* infections in humans do not alter the fecal microbiota [5, 6]. Sometimes the gut microbiota protects its host against helminth infection, such as the maternally transmitted bacterium *Spiroplasma* that protects *Drosophila neotestacea* against nematode parasitism [7], or *Bifdobacterium animalis* that protects mice against *Strongyloides venezuelensis* infection [8]. A recent study demonstrated that *T.muris* infection altered the gut microbiota in mice, and inhibited subsequent rounds of infection [9]. Conversely, *Lactobacillus* facilitates *H. polygyrus* and *T. muris* infections in mice [10, 11]. These opposing findings illustrate the inconsistent nature of helminth-microbiome interactions within their shared hosts. What biological variables explain such heterogeneous results? If we can identify the causes of these variable helminth-microbe associations, then we may be better able to treat dysbiosis and/or macroparasite infections [12–14].

The mechanisms underlying helminth-microbiome interactions are still being elucidated. Macroparasites can interact directly with the microbiome, for instance by secreting antibacterial peptides (e.g., the gastrointestinal nematode *Heligmosomoides polygyrus* secretes at least eight products with antibacterial effects [15]). Or, parasites can indirectly alter the gut microbiota via changes in host physiology, especially immune state. *H. polygyrus bakeri* infection induces colonic regulatory T cells in mice, which are widely regarded to arise in response to gut microbiota colonization [16], and protect mice from colitis [17]. Also, *H. polygyrus* can negatively regulate host intestinal mucosal IL-22 and IL-17, and decrease the host’s expression of epithelial antimicrobial peptides [18]. But non-immune mechanisms also occur: helminth damage to the host gut epithelium can cause malnutrition that changes the microbiome [19].

Indirect interactions between the parasite and microbiota, acting through host traits, should presumably be contingent on host traits. Hosts often vary in their immune response to helminth infection, due to immunogenetic polymorphism [20, 21], energetic reserves [22], stress [23], and prior parasite exposure [24]. Males and females within a species also can respond differently to infections [25], because sex hormones modulate immune traits [26]. But, we lack evidence that variation within host species changes the microbial response to parasitic infection. Here, we show that host sex and autosomal genotype alter the gut microbiota response to cestode infection. We experimentally exposed laboratory-bred threespine stickleback (*Gasterosteus aculeatus*) fish to their native cestode parasite (*Schistocephalus solidus*). We then assayed the gut microbial response to infection and host genotype, and non-additive interactions between infection and genotype. This host-parasite system is especially valuable for such studies because the microbiota have little opportunity for direct interaction with this cestode parasite. After the fish ingests infected copepods, the parasite exits the copepod and rapidly penetrates the intestinal wall (within hours) to establish a long-term infection in the peritoneal cavity, physically separated from the gut microbiota. So, cestode effects on the microbiota are likely to be indirect, mediated via host traits such as immune responses which are often polymorphic within, and divergent between, host species. In an initial exposure experiment (experiment 1), we show that cestode exposure suffices to alter the gut microbiota, even when the infection subsequently fail, but this effect varied among host full-sibling families. In a follow-up experiment (experiment 2) we show that infection presence/absence (controlling for exposure) does also alter the gut microbiota, but this effect varies between host sexes (which are genetically determined) and autosomal genotypes. This result demonstrates that genetic variation in a host species modifies the ecological interaction between helminth and microbiome communities.

## Results

### Effects of cestode exposure versus infection

Experiment 1 was designed to test if exposure alone, or infection presence, change microbial communities. This experimental design is illustrated in Additional file: Figure S1a and sample sizes of fish used in this experiment was listed Additional file: Table S1. The results show that cestode-exposed fish (infected or uninfected) had different microbial communities than the sham-exposed control fish (different unweighted PCoA1 axis scores), though this trend varied among host full-sib families (Fig. 1a and Additional file: Table S2; fish family *P*<0.0001, exposure *P*=0.2589, exposure*family *P*=0.0363). The significant exposure by family interaction occurs because one full-sib family (GG12) showed an atypical microbial response to exposure. Omitting this one family, the remaining families all showed a consistent microbiome response to parasite exposure (exposure *P*=0.0378, family *P*=0.0579, exposure*family *P*=0.1675). In contrast, we observed no difference between infected versus exposed-but-uninfected fish (Fig. 1b and Additional file: Table S2). However, this initial experiment was had modest statistical power, so in Experiment 2 we focused on evaluating only the effect of cestode presence/absence, among fish who were all exposed to the cestode.

**Fig. 1.**
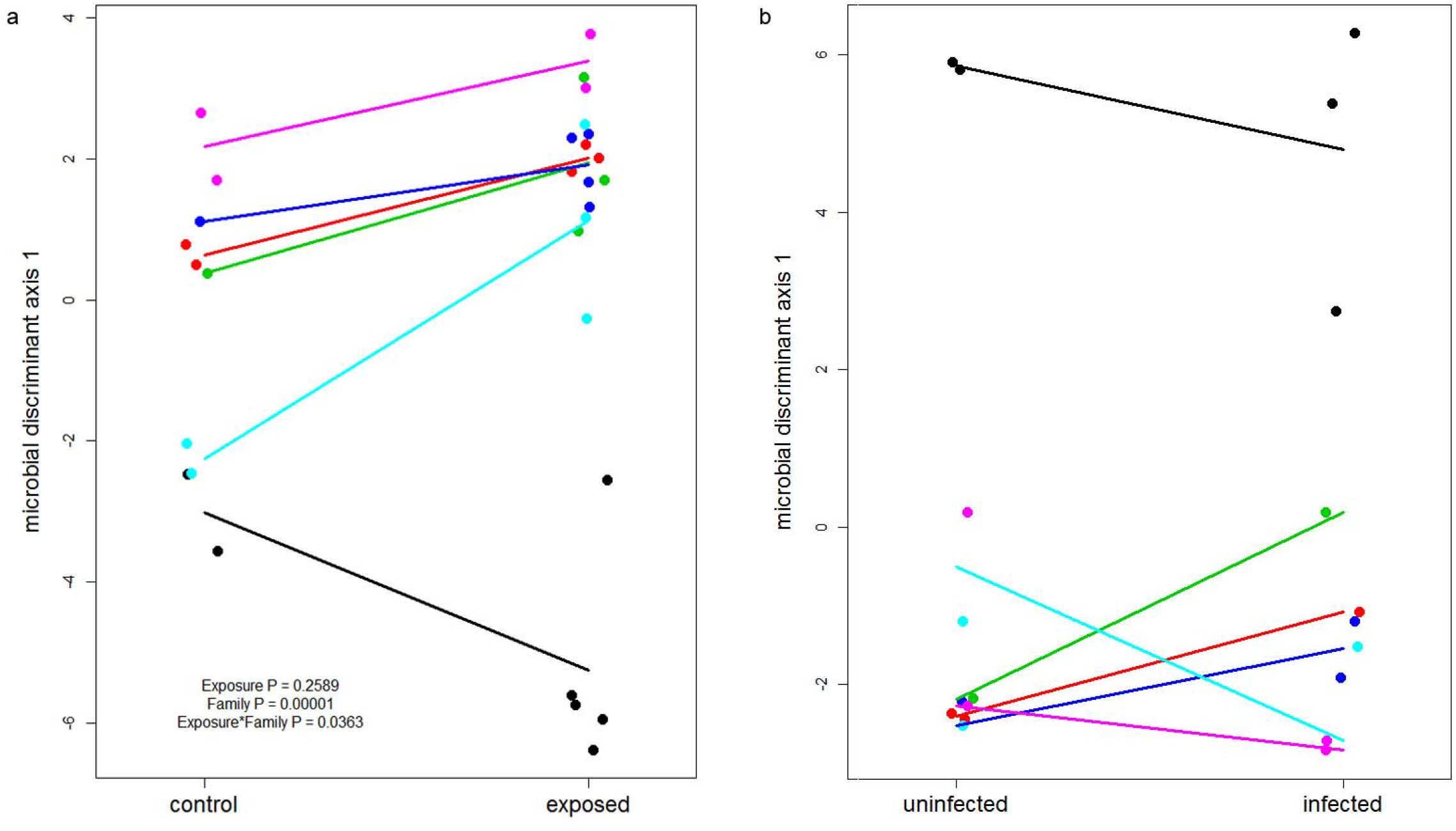
Microbiome community structure of stickleback differs between cestode exposure versus infection. For the purpose of plotting, microbiome community is measured here by scoring fish along the first linear discriminant axis trained on three groups of fish (control, exposed, and infected). The six families are identified by separate point and line colors, underscoring one family with an atypical response to exposure that generated a significant family by exposure interaction; without this family there is a significant exposure main effect. (a) Microbiome community structure differs between stickleback that were experimentally exposed to *S.solidus* (‘exposed’) versus control fish that were sham infected. (b) No significant effect of cestode infection success on experimentally exposed sticklebacks’ gut microbiota.

### Gut microbial community composition

In the Experiment 2, we exposed 711 lab-raised adult F2 hybrid sticklebacks to *S.solidus*, resulting in 256 successful infections (the experimental design is illustrated in Additional file: Figure S1b and sample sizes of this experiment was listed in Additional file: Table S3). We obtained 16S sequences 693 of these 711 fish (a few individuals’ intestines were not retained), yielding 11,586 Operational Taxonomic Units (OTUs). We retained an average of 11,066 sequence reads per fish (s.d.=13,394; median=7,575; range from 48 to 126,943). We excluded all individual stickleback with fewer than 500 sequence reads (*N*=43 removed, 650 retained, Additional file: Table S4). A summary of the gut microbial community composition is provided in Dataset 1. The five most abundant Phyla in the sticklebacks’ intestines were Firmicutes (61.07% of reads on average), Proteobacteria (28.68%), Actinobacteria (5.13%), Bacteroidetes (0.93%) and Planctomycetes (0.67%). The microbial Orders Bacillales (0.53% ∼ 98.34%), Burkholderiales (0.095% ∼ 91.79%) and Actinomycetales (0.027% ∼ 98.46%), and Family Alcaligenaceae (0.025% ∼ 98.27%) were detected in all individual stickleback. As previously noted [27, 28], the stickleback gut microbiota differed dramatically among individuals (Additional file: Figure S2), despite their similar ages and being reared on the same foods in the same research facility (split between two adjacent rooms).

### Host sex, cross, and infection jointly affect the gut microbiome

General linear model analyses of the 181 most common microbial Families (those present in at least 20 fish) suggest that cestode infection and host genotype jointly affect the stickleback gut microbiome. All models included room effect as a covariate to account for the large effect of rearing room (Additional file: Figure S3). Host sex, cross, and mass each had significant main effects on nearly a third of microbial Families (32.6%, 29.8%, and 27.1% respectively; Additional file: Figure S4 and Dataset 2). Main effects of cestode infection were relatively uncommon, however, affecting only 11.0% of microbial Families (still significantly more than expected from our 5% type I error rate; χ^2^=12.7, *P*=0.0003).

Although relatively few microbial Families were associated with cestode presence/absence (consistent with the lower-powered Experiment 1), substantially more Families (50 of 181, or 27.6%) exhibit interactions between infection and other host traits. Specifically, 19.9% of microbial Families exhibited significant infection*cross interactions (contrasting ROB backcross, F2 intercross, or GOS backcross), and 17.1% of Families depend on infection*sex interactions. Fish mass had little effect on the cestode-microbe interaction: only 8.8% of microbial Families exhibited a mass*infection interaction, slightly more common than the 5% false positive rate (χ^2^=4.8, *P*=0.027). Thus, the statistical main effect of infection (our focus in Experiment 1) underestimated the impacts of cestode presence, the majority of which were contingent on genetic characteristics of the host (sex, cross type). Comparable trends were observed if we examined other taxonomic ranks (e.g., 84 common microbial Orders, Fig. 2).

**Fig. 2.**
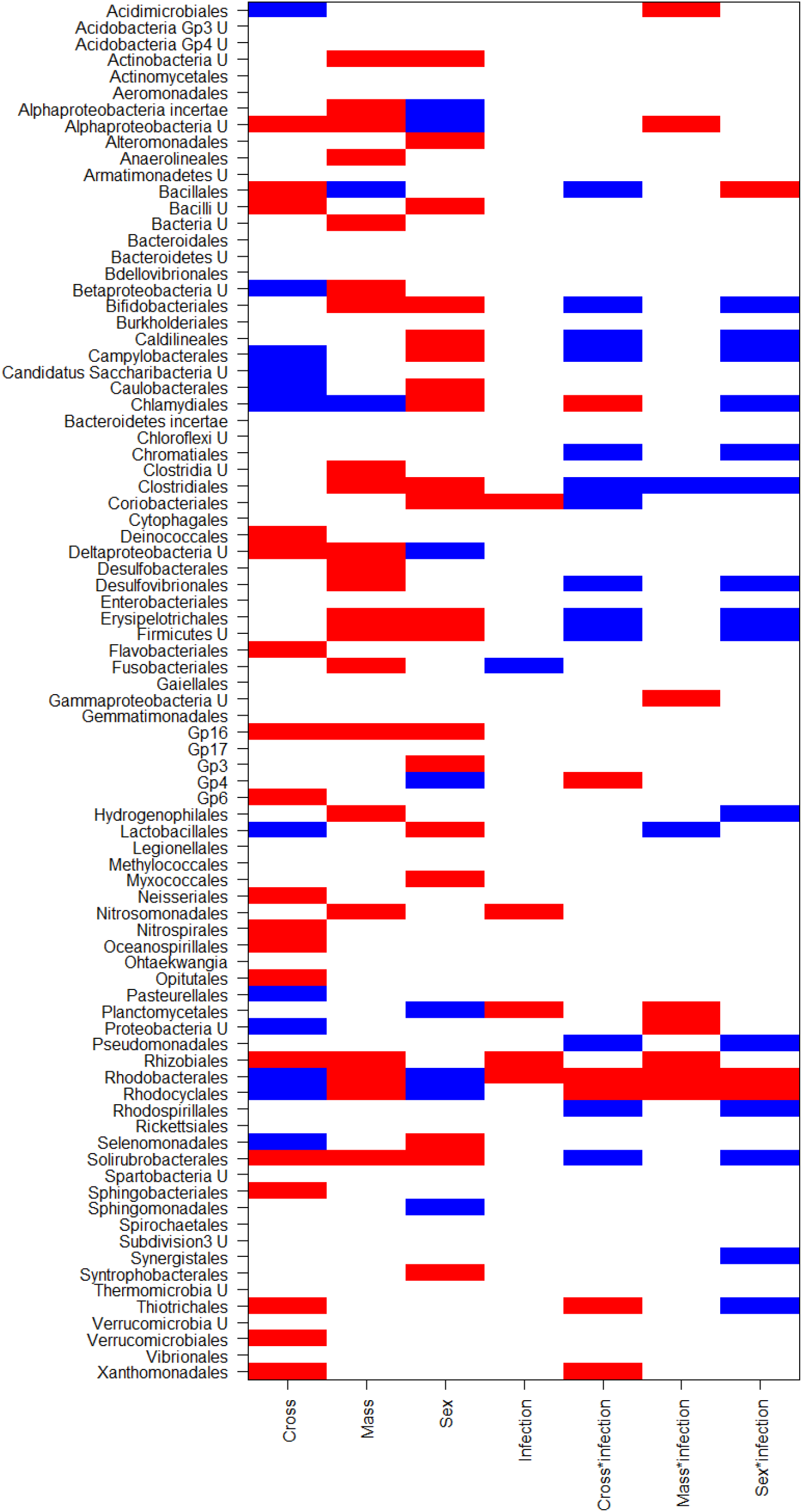
Many microbial Orders (*N*=84, found in a minimum of 20 stickleback) exhibit significant effects of infection or fish characteristics. Here, we identify which microbial Orders exhibit various main and interaction effects. For each Order, a cell is colored if the relevant effect is significant at *P*<0.05. Note that there is a strong excess of significant results at this threshold above the null expectation of a 5% false positive rate (χ^2^ tests *P*<0.0001 for all but the mass*infection interaction). Generalized linear model results for all common Orders (found in at least 20 fish) are summarized in File S2. Color denotes an effect direction. For cross, red denotes higher abundance with more Roberts Lake alleles (ROB backcross), blue denotes higher abundance in Gosling Lake alleles (GOS backcross). For mass, red denotes higher abundance in larger fish, blue in smaller fish. For sex, red denotes higher abundance in males, blue higher in females. For infection, red denotes higher abundance in infected than in uninfected fish, blue is the reverse. For the cross*infection interaction, red indicates instances where infection increases microbe abundance in ROB backcrosses while blue indicates infection increases abundance in GOS. For mass*infection, red indicates that infection increases microbe abundance more strongly in larger fish. For sex*infection, red denotes cases where infection increases microbe abundance most strongly in males, whereas blue implies infection increases microbe abundance mostly in females.

### Examples of interactions between helminth infection and host genotype

A few illustrative examples of such interaction effects are plotted in Fig. 3 and Additional file: Figure S5. We observed a strong main effect of infection for some microbe Orders, such as Fusobacteriales (Fig. 3a and Additional file: Figure S5a, infection effect t=-2.65, *P*=0.0083), which were less abundant in infected sticklebacks’ gut regardless of sex or genotype. As an example of a main effect of host sex, Lactobacillales were more abundant in male than in female stickleback regardless of infection status (Fig. 3b and Additional file: Figure S5b, sex effect t= 3.37, *P*<0.001).

**Fig. 3.**
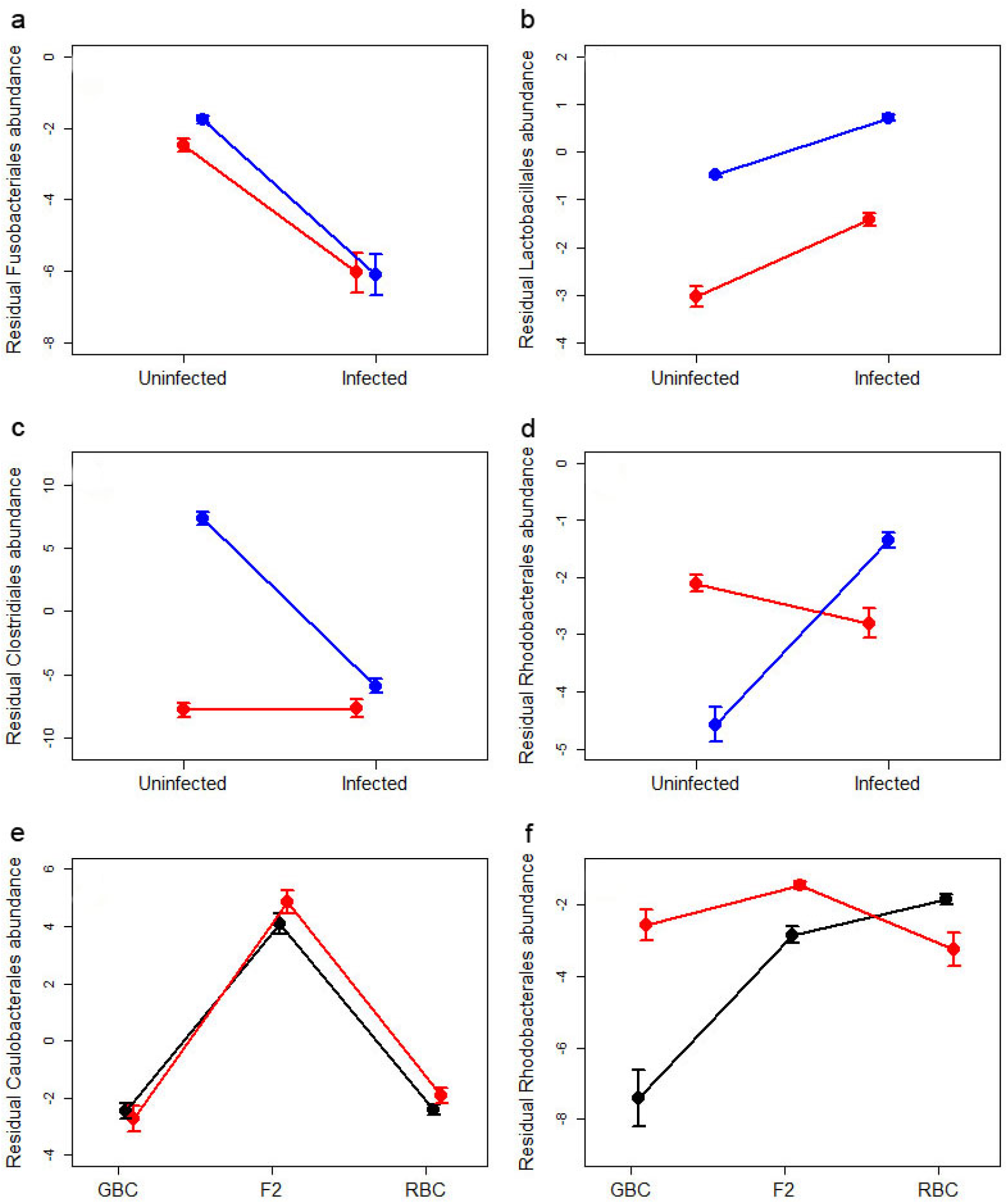
Examples of how the gut microbiome composition depends on host sex, genotype (cross), and infection status, focusing on the relative abundance of microbial Orders. To visualize these effects we first adjusted for variation due to rearing location (room), by calculating residuals from a quasibinomial general linear model of the focal Order’s read counts (out of the total reads per fish) regressed on rearing room. Here we plot the mean (and ±1 s.e. confidence intervals) for this residual abundance, for various groups of fish to illustrate infection and host effects. (a) Fusobacteriales exhibit a significant decrease in cestode-infected fish, both males (blue) and females (red). (b) Lactobacillales are more abundant in males than in females regardless of infection status. (c) Clostridiales abundance is reduced in infected males, but not infected females. (d) Rhodobacterales are more common in females for uninfected fish, but more common in males for infected fish. (e) Caulobacterales are more abundant in F2 hybrids than in either backcross, regardless of infection status (black = uninfected, red = infected). (f) Rhodobacterales are more abundant in fish with a greater fraction of Roberts lake ancestry, but only among uninfected fish; infection leads to a higher Rhodobacterales abundance that is similar across fish genotypes. Fig. S5 shows the same plots, but with all data points included to show the small effect size relative to high among-individual variation.

Clostridiales illustrate the contingent effect of parasite infection (Fig. 3c and Additional file: Figure S5c), with a sex main effect (t=5.56, *P<*0.0001) and a sex*infection interaction (t=-7.46, *P*<0.0001). Clostridiales abundance was insensitive to infection in female hosts, but *S.solidus* infection strongly reduced Clostridiales abundance in male fish. Equivalently, one could say that Clostridiales abundance differed strongly between uninfected males and females, but did not differ between infected males and females. Rhodobacterales also show a sex*infection interaction Fig. 3d and Additional file: Figure S5d, increasing with infection in males but decreasing with infection in females. As a result, among uninfected fish Rhodobacterales were most abundant in females, whereas among infected fish this taxon was most abundant in males.

Host autosomal genotype also influenced the gut microbiota, as indicated by widespread differences between backcross versus F2 hybrids. Caulobacterales were significantly more common in F2 hybrids (Fig. 3e and Additional file: Figure S5e, cross main effect F_2,629_=71.6, *P*<0.0001) than in either backcross. Other Orders were more common in one particular cross, or exhibited an additive trend (F2s intercrosses being intermediate between backcrosses). This autosomal effect altered the impact of cestode infection. Rhodobacterales were more abundant in hosts with greater ancestry from Roberts Lake, but only in the absence of the cestode (Fig. 3f and Additional file: Figure S5f). In contrast, in cestode-infected fish Rhodobacterales were uniformly common in all genotypes.

### Discriminant function analyses of host cross and sex effects on microbiota

We next used multivariate analyses to evaluate the response of the overall gut microbiome community to infection and host genotype (summarized in Additional file: Table S5). LDA separating the four combinations of sex and infection confirmed that helminth infections alter the gut microbiome but sex modifies the effects of helminthiasis (Fig. 4a). The two leading LDA axes, respectively, exhibited a significant effect of sex (F_1,466_=90.8, *P*<0.0001), and an effect of infection (F_1,466_=4.64, *P*=0.0317). For LDA axis 1 (LDA1), these variables also interacted (F_1,466_=9.6, *P*=0.0021). Overall, cestode infection changed female gut microbiota composition more strongly than it changed male microbiota.

**Fig. 4.**
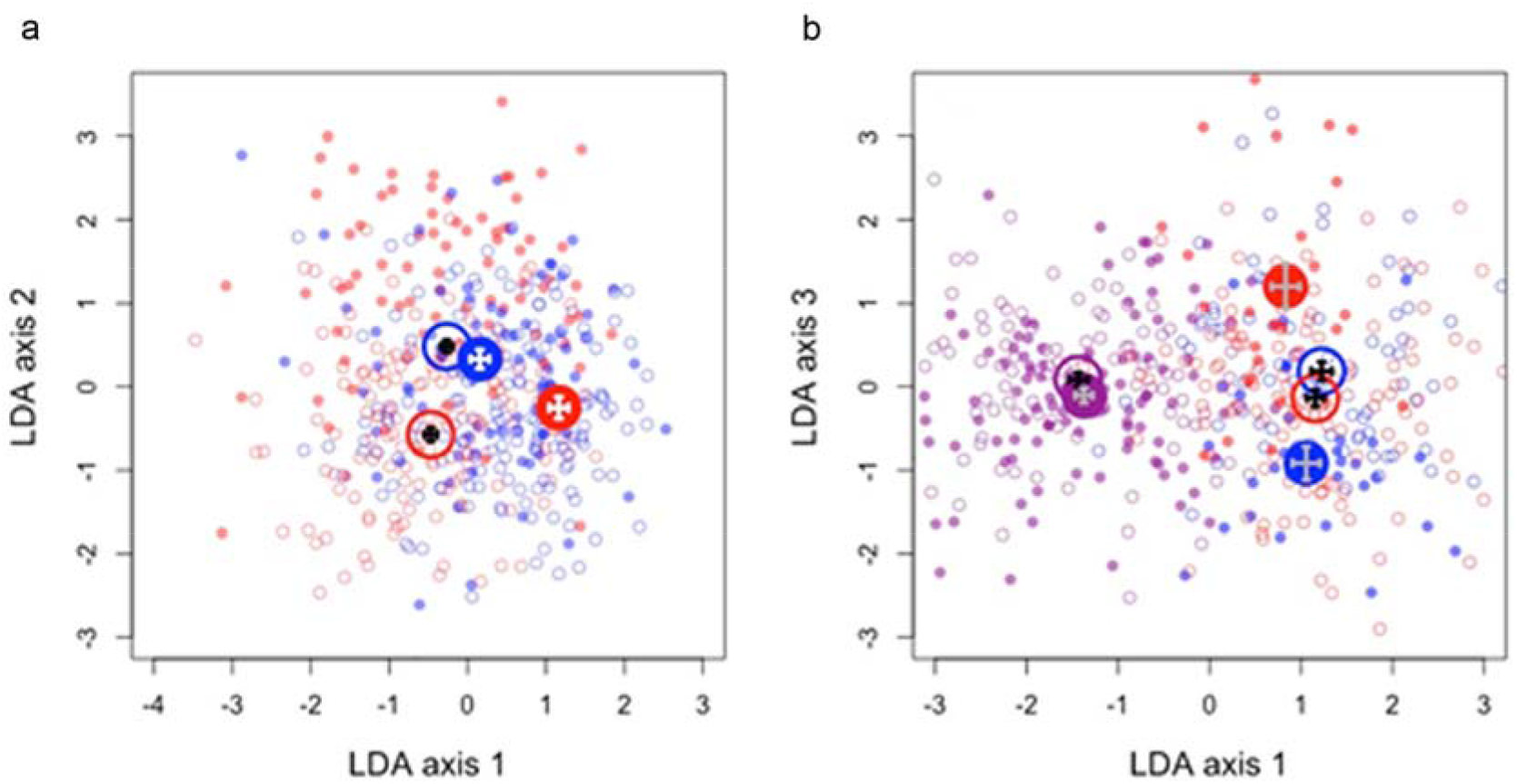
Linear discriminant analysis of the top 50 unweighted PCoA axes of microbial composition reveals effects of host sex, host cross type, and infection status. (a) Results of LDA to separate all combinations of sex and infection status. LDA axis 1 (LDA1) separates infected (filled points) from uninfected (open points; *P*<0.0001). LDA2 separates males and females (blue and red points; *P*<0.0001), and exhibits a significant interaction effect (*P*=0.002). (b) Results of LDA to separate combinations of cross (blue = Gosling backcross, purple = F2, red = Roberts backcross) and infection status (open circles = uninfected, filled circles = infected). LDA1 (68% of variation) separates F2 hybrids from both backcrosses. LDA2 (16% of variation; not shown here) separates Roberts from Gosling backcross fish, with F2 hybrids being intermediate. LDA3 explains 7% of the microbial variation and is most strongly associated with infection status, but in a manner that depends on host cross: the three host crosses are on average almost identical along LDA3 when uninfected (open circles), but diverge when infected with F2s intermediate as expected from additive genetic control. Raw points are shown in faded colors, overlain by larger darker circles representing bivariate averages with ±1 s.e. error bars.

A separate discriminant function analysis of fish cross type and infection status (6 combinations) revealed several insights. The first LDA axis separated F2 hybrid from both reciprocal backcross populations (Fig. 4b), suggesting that there may be transgressive genetic effects on the microbiota (e.g., Fig. 3e). F2 hybrids were intermediate between the backcrosses along LDA2, consistent with additive genetic control of other aspects of the microbiota (Additional file: Figure S6). The third axis separated infected and uninfected fish, but in a highly genotype-dependent manner (Fig. 4b and Additional file: Figure S6). In F2 cross fish, LDA3 scores were insensitive to infection, whereas in both backcross populations the LDA3 scores changed strongly in response to infection. All three crosses had similar LDA3 scores when uninfected, but Gosling and Roberts backcross fishes’ microbiomes diverged in response to infection.

### QTL mapping of gut microbiota composition

Based on the effects of cross type, described above, we expected to be able to locate host loci that (i) explain variation in gut microbiota composition, and (ii) do so differently for infected versus uninfected fish. We used quantitative trait locus (QTL) mapping to test this expectation, and to identify chromosomal regions for future detailed mapping, and possible candidate gene identification and validation.

ddRADseq identified 234 SNPs with fixed differences between the Roberts and Gosling populations and sufficiently deep coverage within and among the F2 hybrid individuals, yielding approximately 10 markers per linkage group. Another paper will describe QTL mapping of infection outcomes and host immune traits (Weber et al, in preparation); here we focus only on mapping the gut microbiota. We located no significant QTL for any of the top 10 weighted PCoA axes, and no significant QTL for microbial diversity (at a rarefaction of 2000 or 4000 reads per fish). But, we did detect weak-effect QTL for an unweighted PCoA axis. Unweighted PCoA axis 5 exhibited three QTLs that narrowly exceeded our stringent threshold for significance (Additional file: Figure S7a-c). We had a stronger signal when we fused the many PCoA axes into a single metric using linear discriminant analysis (trained to distinguish the two backcrosses then applied to all samples). Using this first LDA axis we detected a single well-supported QTL on Chr9 (Additional file: Figure S7d-e). Note that because QTL mapping was done within each cross, using LDA to define an axis that distinguishes between crosses is not tautological. Chromosome 9 does not have any noteworthy effect on cestode infection success (infection QTL described in Weber et al, in preparation).

The lack of strong QTL for microbial ordination metrics led us to hypothesize that host control of the whole microbial community is highly polygenic. If host genetic variation acts on particular microbe taxa, it might act only weakly on PCoA scores and be correspondingly hard to map. So we next mapped microbial Orders separately, revealing numerous taxon-specific QTL. To illustrate, Fusobacterales exhibit two strong autosomal QTL, plus an association with the X chromosome (ChrX) indicating a sex effect (Fig. 5). Other examples are plotted in supplementary figures (Additional file: Figure S8), and their overlapping distributions in the genome are presented in Fig. 6. This summary reveals genomic ‘hotspots’ for QTL affecting the microbiota, on Chr1, Chr2, and Chr3. Many of these microbial Orders also mapped to the sex chromosome (Chr19), consistent with the common main effect of sex. Similar results were obtained for Family level QTL mapping.

**Fig. 5.**
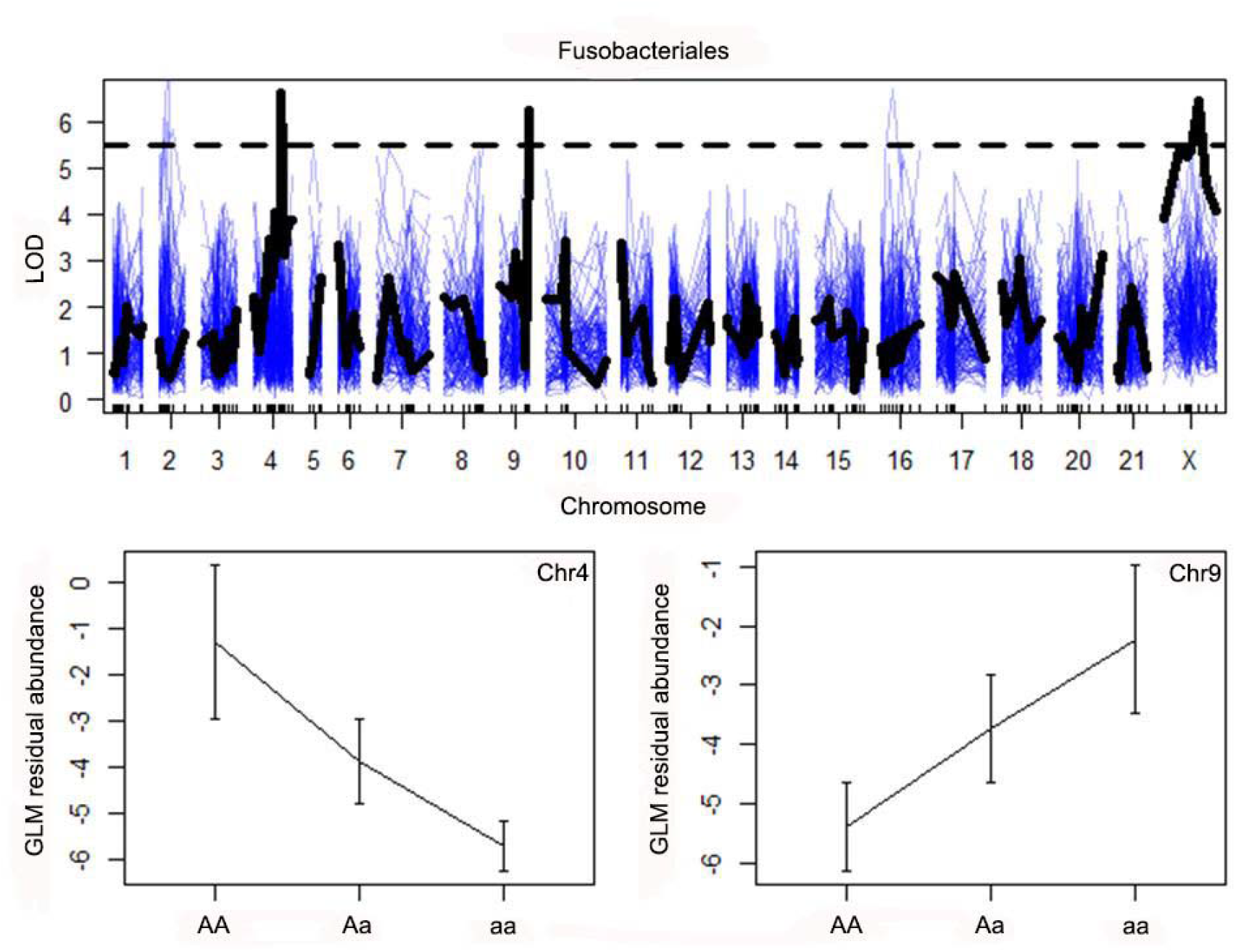
QTL map of Fusobacteriales relative abundance (with non-parametric statistical tests) reveals two autosomal QTL plus an association with the X chromosome. Analyses were run separately for each cross, with rearing room as an additive covariate, then the LOD scores from the three maps were summed. The observed summed LOD scores are plotted in black for each linkage group, measuring statistical association between the focal trait and the chromosomal region. Marker locations are indicated as tick marks along the horizontal axis. Thin blue lines represent null summed LOD scores from within-cross permutations of traits. The horizontal dashed line indicates the upper 99.99% quantile for the null LOD scores. Three QTL exceed this threshold, on chromsomes 4, 9, and X. Two of these are plotted in the lower panels (left, locus X109 on Chr 4, genotype *P*=0.0013; right, locus X156 on Chr 9, genotype *P*=0. 0060). The y axis in these effect plots are the residuals from a regression of unweighted PCoA5 on rearing room. We plot the means microbial abundance with ±1 standard error bars, for each of the three genotypes at each locus.

**Fig. 6.**
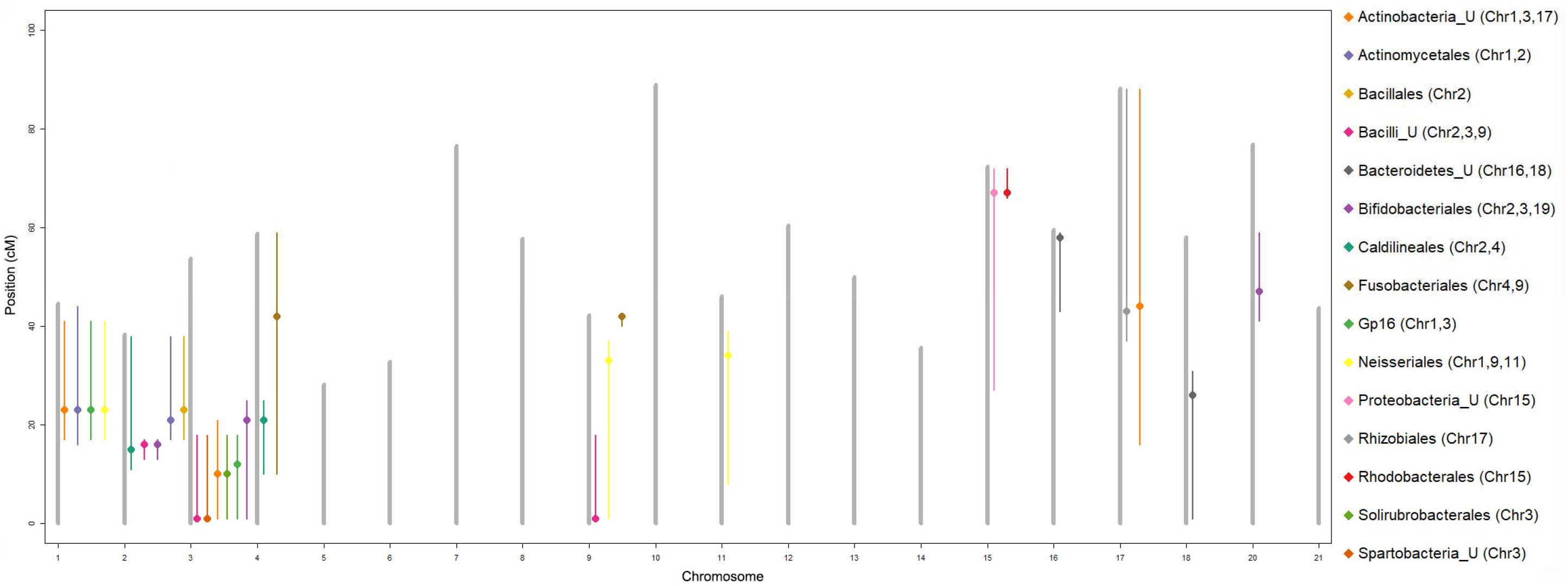
A summary of the locations of QTL affecting the relative abundance of individual microbial Order. Each chromosome is plotted as a vertical grey bar, and to its right we plot microbial QTL located on that chromosome. A dot symbol indicates the location of maximum LOD score for each QTL, and thin vertical lines indicate the inferred width of the QTL. For sex-linked microbes, their QTL span the entire X chromosome, so we omit that linkage group. The key to the right lists the locations of the significant QTL for each focal taxon.

We next tested whether these QTL are contingent on cestode presence or absence, as expected from the interactive effect of infection and cross type, described above. We mapped QTL separately for infected fish and for uninfected fish, and then looked for loci with QTL in only one of these groups. We found numerous host loci that are only associated with microbial variation in uninfected fish, whereas the same locus is unrelated to the microbiota among infected fish (e.g., Additional file: Figure S9a-f). In fewer instances, we identified host QTL for microbiota in infected fish only (Additional file: Figure S9g-h). In several cases stratifying by cestode infection revealed QTL that would not have been detected otherwise. For example, Solirubrobacterales (Additional file: Figure S9e-f) have a QTL on Chr3 that is observed in all fish, but which in fact only acts on the uninfected fish (a larger sample size). When we ignored infection status, the same microbial taxon had a non-significant QTL on stickleback Chr16. The lack of Solirubrobacterales in infected fish was masking the effect of host genetic variation at that site.

## Discussion

The interaction between hosts, parasites, and gut microbes represents a rich opportunity for experimental studies of multi-way ecological interactions. A growing literature suggests that helminth infection can alter the vertebrate gut microbiota [1]. Our experiment adds an important twist to this literature: the magnitude and direction of helminth-induced microbial changes depend on both the sex of the host, and the host’s autosomal genotype. By examining F2 hybrids between two recently-diverged host populations, we were able to identify autosomal loci (QTL) that contribute to variation in microbiota composition, as well as variation in microbiota response to infection. The implication is that genetic variation within one species (the host) alters the ecological effect of another species (the cestode parasite) on a third party (the microbial community).

The inconvenient implication of this finding is that we may not readily generalize helminth-microbe effects beyond the genotypes (and sex) we study. This unwelcome news is tempered by the opportunity it presents: host genetic variation can therefore help us identify the mechanisms by which helminths alter the gut microbiota (or, vice versa). Doing so may allow us to develop a more general predictive model that can account for such heterogeneity among study subjects, and ultimately explain why individual hosts differ in their microbial response to infection.

To date, many studies have evaluated effects of helminth infection on the gut microbiota of animals [1, 13]. Yet, we remain largely ignorant as to whether these helminth effects depend on other biological variables, whether environmental effects or host genotype. Here, we have provided evidence of such interactions: host genotype (sex and autosomal) and helminth infection synergistically alter the microbiota. One third of the common microbial Orders exhibited significant interactions between infection and host genotype (Fig. 2), whereas infection by itself affected only 7% of the Orders. For example, the abundances of some microbial Orders in female hosts were significantly different with those males when uninfected, but no difference between them when infected. Discriminant analysis suggested that females’ microbiota were generally more sensitive to infection than males’ were. This sex-dependent effect of infection is consistent with other recent papers on stickleback suggesting that sex modifies the gut microbiome’s response to diet [27], and MHC genotype [29].

Host autosomal genotype also affected gut microbial composition, and modified microbes’ responses to helminth infection. This inference is consistent with many other studies of vertebrates that have yielded many examples of host genetic control of microbiome composition [30, 31]. What makes the above results novel is that this host genetic variation also alters the microbiome’s response to a third (parasitic) species. We coarsely localize these autosomal effects to a modest number of chromosomal loci (QTL). These QTL do not contain MHCIIb, which has previously been associated with natural variation in stickleback gut microbiota [29]. At present we lack the resolution fine-map down to specific candidate genes, but some intriguing possibilities exist. The QTL on Chr2, which affect multiple microbial Orders, contains transcription factor c-MAF, which activates the expression of IL-4 in Th2 cells and attenuates Th1 differentiation [32]. Interestingly, recently Xu et al. reported c-MAF-dependent regulatory T cells mediate immunological tolerance to intestinal microbiota [33]. In addition, the QTL on this chromosome contains reticulocalbin and synaptotagmin IX, which are associated with calcium ion binding; dietary calcium can affect the intestinal microbiota composition [34–36]. The QTL on Chr3 contains cadherins, which play a key role in intestinal homeostasis and barrier function [37]. Substantial future work is needed to fine-map candidate genes and experimentally validate their suspected effects on the gut microbiota, and on the cestode-microbiome interaction.

Interactions between host genotype and infection likely arise from host genetic variation in immune response to cestodes. The presence of helminths can interfere with TLR or downstream signaling pathways [38], which may have ancillary effects on the microbiota. Adaptive immune responses may be involved as well. Helminths can trigger a Th2 response that increases mucus production and epithelial cell turnover, altering the mucosal microbiota [39]. Any polymorphism in immune genes (e.g. TLRs) involved in these pathways may lead to an exaggerated (or, reduced) immune response to helminth infection, and thereby modify the effect of helminths on gut microbes. This logic applies equally to immune differences between males and females [40, 41].

In the specific case examined here, helminth effects on the microbiota are especially likely to involve indirect effects via host immunity, rather than a direct microbe-helminth interaction. This is because *S.solidus* is not an intestinal parasite. After being ingested, it quickly transits into the body cavity where it persists for months but cannot directly affect the microbiota. Prior studies confirm that stickleback vary in their ability to detect *S.solidus*, mount an effective immune response, and susceptibility to cestode immune suppression [42–44]. Polymorphism in these immune responses may cascade throughout the host body, imposing collateral effects on the microbiota.

The above results specifically apply to variation among cestode-exposed stickleback. But, our Experiment 1 suggests that stickleback gut microbiota depended most strongly on parasite exposure (regardless of outcome), rather than infection itself. Apparently there is a lasting effect of the transient presence of a tapeworm in the stickleback gut lumen, which outweighs the effect of tapeworm presence or absence in the peritoneum. If indeed brief parasite exposure sends lasting ripples through the microbial community, then the study of the wild microbiome will be still more difficult. We rarely if ever know our study animals’ history of unsuccessful infections, so these ripples might generate substantial and untraceable variation among wild individuals. Yet infection success also alters the gut microbiome, as we then revealed in the follow-up Experiment 2.

At present we do not know the fitness consequences of the altered microbiota that we document here. Some studies have reported detrimental helminth effects on the microbiota. Experimental *T. muris* infection in mice alters the microbiota composition leading to reduced availability of microbial metabolomics products needed by the host (vitamin D2/D3 derivatives, many fatty acids, and amino acid synthesis intermediates) [45]. Conversely, helminth infection can be beneficial, such as reducing the risk of *H. pylori*-induced gastric lesions [46]. Our results suggest that these positive or negative effects of helminth-microbiota interactions will be contingent on the specific sex and genotype of the hosts considered.

The absence of helminths in wealthy counties has been postulated to contribute to the increasing prevalence of allergic and autoimmune diseases [47], which might be attributed, at least in part, to alterations to the intestinal microbiota [13]. Some of these auto-immune diseases are sex-biased in humans [48, 49]. Our results raise the possibility that this sex-bias may be related to sex-specific microbial responses to the presence (or, absence) of helminthes. Therefore, we propose that sex-specific environmental effects on the gut microbiota might contribute to understanding sex-specific inflammatory diseases and sex differences in dysbiosis. Recently, helminth-based therapeutics, e.g. infection with *T. suis*, have been tested as possible treatments for inflammatory bowel diseases [47]. If our results hold beyond stickleback, we predict that clinical therapeutic efficacy may differ between sexes, or between host genotypes. Consequently, such treatments might need to take account of these interactions, potentially requiring therapies tailored to host sex or genotype.

## Materials and Methods

### Stickleback breeding

Gosling Lake and Roberts Lake on Vancouver Island, British Columbia contain stickleback populations that differ in immune phenotypes, immune response to infection, parasite growth in the lab, and cestode prevalence in the wild (>70% and 0% respectively) [50, 51]. We generated pure Roberts (ROB), pure Gosling (GOS), and reciprocal F1 hybrid families (RG or GR), and raised these to maturity in the lab, at which time we interbred them to generate second generation (F2) hybrids. These F2 hybrids included intercross families (F1*F1 matings) and reciprocal backcross families (ROB*F1, F1*ROB, GOS*F1, F1*GOS). All fish were reared in freshwater conditions and housed with full-siblings (9-39 animals per family) in either 40-L or 10-L tanks.

### Parasite Collection and Experimental Exposures

These fish crosses and infections were designed to obtain a quantitative trait locus (QTL) map of the genetic basis of host control of cestode growth and viability; those results will be reported elsewhere (Weber et al, in preparation). Here, we focus on testing the hypothesis that host genetic variation alters the impact of helminth infection on the vertebrate gut microbiota. We conducted two sets of experimental infections (Additional file: Figure S1).

(Experiment 1) As a pilot study, we evaluated the effects of cestode exposure, and successful infection, on the host gut microbiome. We experimentally exposed adult pure-bred fish from six full-sib families from Gosling Lake, to *S.solidus*. Within each family, five individuals were controls fed uninfected copepods (sham exposure), while the rest received infected copepods, only some of whom were ultimately infected. This experimental design yielded three categories of fish within each family: unexposed controls, exposed-but-uninfected controls, and infected fish (*N*=30, Additional file: Figure S1a and Table S1). For these exposures we followed a standard procedure described by Weber et al. [50]. Briefly, we dissected mature *S.solidus* out of infected wild-caught fish from Gosling, and bred the tapeworms in culture media in dark waterbath (mimicking the gut of piscivorous birds), to collect eggs. (Roberts Lake lacks *S.solidus*, so this cannot be used as a source population.) We hatched the tapeworm eggs and fed them to copepods (*Macrocyclops albidus*), then visually isolated infected copepods to feed in controlled doses to the lab-raised stickleback. Forty-three days after parasite exposure, we euthanized fish with MS-222 to check for infection success. We froze the entire intestine in ethanol for subsequent microbiome analysis.

(Experiment 2) To evaluate the effect of host genotype on cestode-microbiome interactions, we experimentally exposed adult F2 hybrid fish (intercross and backcross, *N*=711, Additional file: Figure S1b and Table S3) to *S.solidus*, then assayed infection outcomes and sequenced the gut microbiome. All fish from this experiment were exposed to *S.solidus*, but not all were infected successfully, providing a contrast between fish with versus without the parasite, controlling for initial exposure history. By excluding the unexposed (sham) treatment, we are able to devote more statistical power to evaluating host genotype interactions with cestode presence versus absence, without the substantial additional confounding variation associated with parasite exposure (see Experiment 1 results).

### Amplification, sequencing and analysis of 16S rRNA amplicons

We extracted DNA from the entire intestine (both content and mucosa) of stickleback (*N*=30 pure-bred GOS fish in Experiment 1; *N*=693 F2 hybrids in Experiment 2) using the MoBio Powersoil DNA Isolation Kits, as described by Bolnick et al. [27, 28]. For the pure-bred GOS fish, we used the V4-V5 primers [52]. For the F2 fish, 16S rRNA amplicons were generated for the V4 hypervariable [27]. Sample-specific barcodes were used as described in Bolnick et al. [27]. PCR amplification was performed in triplicate for each sample in a reaction volume of 25 µL containing 1×Q5 High-Fidelity Master Mix, 5 pmol forward and reverse primers and 1µL template DNA. The PCR products for each sample were pooled and quantified with Picogreen double-stranded DNA reagent to facilitate pooling equimolar amounts of amplicons for sequencing. To test for contamination, negative controls (without samples) were set up for both the DNA extraction and 16S PCR amplification stages. The results indicated there was no detectable contamination because the PCRs yielded negligible DNA concentrations during Picogreen quantitation. Amplicon pools were paired-end sequenced on an IlluminaMiseq platform at GSAF (Genomic Sequencing and Analysis Facility) at the University of Texas at Austin.

The raw paired-end reads were demultiplexed, and subsequent sequence processing was performed using the mothur software package (v.1.39.1), following standard operating procedures (SOP) [53, 54] (https://www.mothur.org/wiki/MiSeq_SOP). Briefly, the sequences were trimmed for 16S rRNA gene primer sequences, and then assembled into contigs and aligned with 16S rRNA gene sequences from the ARB Silva v128 reference database [55]. Chimeric sequences were detected using VSEARCH within mothur in each sample and removed. The remaining sequences were classified by using Bayesian classifier with a training set (version 16) from the Ribosomal Database Project (http://rdp.cme.msu.edu) [56]. Operational Taxonomic Units (OTUs) were identified using the UCLUST algorithm based on 97% similarity. We rarefied the data to 4000 sequences per sample to calculate unweighted and weighted UniFrac distance and PCoA scores by phyloseq library in R [57].

### Analyses of pure GOS fish experimentally exposed to *S.solidus* or a negative control

We sequenced the gut microbiota of GOS fish (Experiment 1) to obtain counts of microbe abundance. We used mixed model linear models to analyze effects of family and infection status on unweighted or weighted PCoA axis 1. Family was a random effect, with a random family*infection interaction. First, we contrasted unexposed (sham infections) versus exposed fish (regardless of the outcome of exposure). Second, we compared uninfected fish (sham or failed infections) versus infected fish.

### Analyses of F2 hybrid fish experimentally exposed to *S.solidus*

#### General Linear Model Analyses

Evaluating the results of Experiment 2, we used quasibinomial general linear models (GLMs), implemented in R, to examine the effects of infection and host genotype on microbial composition (the relative abundance of each commonly observed Order/Family, found in at least 20 fish [*N* = 84 Orders/181 Families]). For each taxon, we estimated a GLM in which the fraction of reads attributed to that taxon (out of all reads) was related to *S.solidus* presence/absence, and effects of host cross (Gosling backcross, F2 intercross, or Roberts backcross), host sex, and host mass. We included interactions of particular interest to us here: infection*cross, infection*sex, and infection*mass. As fish were reared in two rooms, we used a room effect as a covariate. We used a sequential Bonferroni correction to P-values when evaluating effects of particular microbial Orders/Families.

#### Discriminant Function Analyses

We next examined the whole-microbiome effects of infection, sex, and cross direction. We applied linear discriminant analysis (LDA) to the top 50 weighted/unweighted microbial PCoA axes (which cumulatively account for 99.99% of variance in microbial alpha diversity using phylogenetically weighted or unweighted presence-absence data). We first used LDA to distinguish four groups (combinations of sex and infection status). Second, we used LDA to distinguish six groups (factorial combinations of three host cross types and infection status). We used ANOVAs to test for effects of sex, cross, and infection status on each LDA axis. We also confirmed these statistical effects using a MANOVA applied directly to the top 50 PCoA axes to test for effects of sex, cross, infection, host mass, and interactions among these variables.

#### QTL mapping: ddRAD genotyping

We used a Promega Wizard SV 96-well plate kit to extract DNA from alcohol-preserved fin clips from all F2 hybrid fish (intercross and backcrosses), as well as all grandparents used to generate the crosses. We then genotyped the fish to obtain SNPs for quantitative trait locus (QTL) mapping, using the ddRADseq protocol described in Peterson et al. [58], with bioinformatics steps to identify SNPs conducted as described in Stuart et al. [59]. We retained only SNPs exhibiting fixed differences between the Roberts and Gosling Lake grandparents (e.g., fully informative for QTL mapping). This conservative approach yielded 236 genetic markers for mapping, on average slightly more than 10 markers per linkage group.

#### QTL mapping: analyses

We mapped quantitative trait loci (QTL) for several microbiome measures: alpha diversity (using 2000 or 4000 read depth normalization), the top 10 weighted PCoA axes (or unweighted axes), and the relative abundance of the common microbial Orders (as described above for GLMs). We built linkage maps for each cross separately in R/qtl [60], and used the *scanone* function with ‘hk’ interval mapping (using rank-based nonparametric tests for microbial Order relative abundance). To account for the complex cross design (with backcrosses and F2 intercrosses), we built separate QTL maps within each of the three cross types, then summed their LOD scores. This yields one summary statistic measuring a locus’ association with the focal trait, while accounting for between-cross differences in QTL effect size or marker linkage. We compared this summed LOD against null expectations obtained by 1,000 permutations of the focal phenotype across fish within each cross, each time redoing each cross’ QTL map and summing cross null LOD scores. Conservatively, we consider an observed QTL significant when its summed LOD exceeded the 99.99% quantile from that marker’s null values at that same marker. We double-checked each significant QTL with a GLM (as described above) testing for fish genotype effect (at the nearest genetic marker) on the focal microbiome variable.

## Acknowledgements

We thank the Genome Sequencing and Analysis Facility (GSAF) at the University of Texas at Austin for sequencing support, and the Texas Advanced Computing Center (TACC) and High Performance Computing platform of Northwest A&F University for computational resources. This research was supported by funding from NSFC (Grant 31672680) to Fei Ling, and the Howard Hughes Medical Institute to DIB, NIH (Grant 1R01AI123659-01A1) to DIB.

## Authors’ contributions

D.I.B., J.W., and N.C.S. designed the study; F.L., N.C.S., and L.M. performed the study; D.I.B., F.L., C.S., D.C., B.Z., and G.X.W. analyzed the data; D.I.B., and F.L wrote the paper.

## Availability of data materials

Sequence data have been deposited in the Sequence Read Database (SRA) under project IDs SRP115642 (BioProject PRJNA398629) and SRP115678 (BioProject PRJNA398630) for experimental 1 and 2, respectively. All other relevant data are available in this article and its Supplementary Information files, or from the corresponding authors upon request.

## References

1. Reynolds LA, Finlay BB, Maizels RM. Cohabitation in the intestine: Interactions among helminth parasites, bacterial microbiota, and host immunity. J Immunol. 2015;195:4059–66.

2. Peachey LE, Jenkins TP, Cantacessi C. This gut ain’t big enough for both of us. Or is it? Helminth-microbiota interactions in veterinary species. Trends Parasitol. 2017;3:619–32.

3. McKenney EA, Williamson L, Yoder AD, Rawls JF, Bilbo SD, Parker W. Alteration of the rat cecal microbiome during colonization with the helminth *Hymenolepis diminuta*. Gut Microbes. 2015;6:182–93.

4. Aivelo T, Norberg A. Parasite-microbiota interactions potentially affect intestinal communities in wild mammals. J Anim Ecol. 2018;87:438–47.

5. Cooper P, Walker AW, Reyes J, Chico M, Salter SJ, Vaca M, Parkhill J. Patent human infections with the whipworm, *Trichuris trichiura*, are not associated with alterations in the faecal microbiota. PLoS One. 2013;8:e76573.

6. Cantacessi C, Giacomin P, Croese J, Zakrzewski M, Sotillo J, McCann L, Nolan MJ, Mitreva M, Krause L, Loukas A. Impact of experimental hookworm infection on the human gut microbiota. J Infect Dis. 2014;210:1431–4.

7. Jaenike J, Unckless R, Cockburn SN, Boelio LM, Perlman SJ. Adaptation via symbiosis: recent spread of a *Drosophila* defensive symbiont. Science. 2010;329:212–5.

8. Oliveira-Sequeira TC, David ÉB, Ribeiro C, Guimarães S, Masseno AP, Katagiri S, Sequeira JL. Effect of *Bifidobacterium animalis* on mice infected with *Strongyloides venezuelensis*. Rev Inst Med Trop Sao Paulo. 2014;56:105–9.

9. White EC, Houlden A, Bancroft AJ, Hayes KS, Goldrick M, Grencis RK, Roberts IS. Manipulation of host and parasite microbiotas: Survival strategies during chronic nematode infection. SciAdv. 2018;4:eaap7399.

10. Dea-Ayuela MA, Rama-Iniguez S, Bolas-Fernandez F. Enhanced susceptibility to *Trichuris muris* infection of B10Br mice treated with the probiotic *Lactobacillus casei*. Int Immunopharmacol. 2008;8:28–35.

11. Reynolds LA, Smith KA, Filbey KJ, Harcus Y, Hewitson JP, Redpath SA, Valdez Y, Yebra MJ, Finlay BB, Maizels RM. Commensal-pathogen interactions in the intestinal tract: Lactobacilli promote infection with, and are promoted by, helminth parasites. Gut Microbes. 2014;5:522–32.

12. Loke P, Lim YA. Helminths and the microbiota: parts of the hygiene hypothesis. Parasite Immunol. 2015;37:314–23.

13. Zaiss MM, Harris NL. Interactions between the intestinal microbiome and helminth parasites. Parasite Immunol. 2016;38:5–11.

14. Rapin A, Harris NL. Helminth-bacterial interactions: Cause and consequence. Trends Immunol. 2018;39:724–33.

15. Hewitson JP, Harcus Y, Murray J, van Agtmaal M, Filbey KJ, Grainger JR, Bridgett S, Blaxter ML, Ashton PD, Ashford DA, Curwen RS, Wilson RA, Dowle AA, Maizels RM. Proteomic analysis of secretory products from the model gastrointestinal nematode *Heligmosomoides polygyrus* reveals dominance of venom allergen-like (VAL) proteins. J Proteomics. 2011;74: 1573–94.

16. Geuking MB, Cahenzli J, Lawson MA, Ng DC, Slack E, Hapfelmeier S, McCoy KD, Macpherson AJ. Intestinal bacterial colonization induces mutualistic regulatory T cell responses. Immunity 2011;34:794–806.

17. Hang L, Blum AM, Setiawan T, Urban JP Jr, Stoyanoff KM, Weinstock JV. *Heligmosomoides polygyrus bakeri* infection activates colonic Foxp*^3+^* T cells enhancing their capacity to prevent colitis. J Immunol. 2013;191:1927–34.

18. Su L, Su CW, Qi Y, Yang G, Zhang M, Cherayil BJ, Zhang X, Shi HN. Coinfection with an intestinal helminth impairs host innate immunity against *Salmonella enterica* serovar Typhimurium and exacerbates intestinal inflammation in mice. Infect Immun. 2014;82:3855–66.

19. Glendinning L, Nausch N, Free A, Taylor DW, Mutapi F. The microbiota and helminths: sharing the same niche in the human host. Parasitology. 2014;141:1255–71.

20. Grant AV, Araujo MI, Ponte EV, Oliveira RR, Cruz AA, Barnes KC, Beaty TH. Polymorphisms in IL10 are associated with total Immunoglobulin E levels and *Schistosoma mansoni* infection intensity in a Brazilian population. Genes Immun. 2011;12:46–50.

21. Costa RD, Figueiredo CA, Barreto ML, Alcantara-Neves NM, Rodrigues LC, Cruz AA, Vergara C, Rafaels N, Foster C, Potee J, Campbell M, Mathias RA, Barnes KC. Effect of polymorphisms on TGFB1 on allergic asthma and helminth infection in an African admixed population. Ann Allergy Asthma Immunol. 2017;118:483–8.e1.

22. Janet K, Tommy LFL. Flying with diverse passengers: greater richness of parasitic nematodes in migratory birds. Oikos. 2015;124:399–405.

23. Guivier E, Bellenger J, Sorci G, Faivre B. Helminth interaction with the host immune system: Short-term benefits and costs in relation to the infectious environment. Am Nat. 2016;188:253–63.

24. Bourke CD, Maizels RM, Mutapi F. Acquired immune heterogeneity and its sources in human helminth infection. Parasitology. 2011;138:139–59.

25. Fischer J, Jung N, Robinson N, Lehmann C. Sex differences in immune responses to infectious diseases. Infection. 2015;43:399–403.

26. Klein SL. The effects of hormones on sex differences in infection: from genes to behavior. Neurosci. Biobehav. Rev. 2000,24:627–38.

27. Bolnick DI, Snowberg LK, Hirsch PE, Lauber CL, Org E, Parks B, Lusis AJ, Knight R, Caporaso JG, Svanbäck R. Individual diet has sex-dependent effects on vertebrate gut microbiota. Nat Commun. 2014;5: 4500.

28. Bolnick DI, Snowberg LK, Hirsch PE, Lauber CL, Knight R, Caporaso JG, Svanbäck R. Individuals’ diet diversity influences gut microbial diversity in two freshwater fish (threespine stickleback and Eurasian perch). Ecol Lett. 2014;17:979–87.

29. Bolnick DI, Snowberg LK, Caporaso JG, Lauber C, Knight R, Stutz WE. Major Histocompatibility Complex class IIb polymorphism influences gut microbiota composition and diversity. Mol Ecol. 2014;23:4831–45.

30. Benson AK, Kelly SA, Legge R, Ma F, Low SJ, Kim J, Zhang M, Oh PL, Nehrenberg D, Hua K, Kachman SD, Moriyama EN, Walter J, Peterson DA, Pomp D. Individuality in gut microbiota composition is a complex polygenic trait shaped by multiple environmental and host genetic factors. Proc Natl Acad Sci USA. 2010;07: 18933–8.

31. Goodrich JK, Davenport ER, Waters JL, Clark AG, Ley RE. Cross-species comparisons of host genetic associations with the microbiome. Science. 2016;52:532–5.

32. Ho IC, Lo D, Glimcher LH. c-maf promotes T helper cell type 2 (Th2) and attenuates Th1 differentiation by both interleukin 4-dependent and -independent mechanisms. J Exp Med. 1998;188:1859–66.

33. Xu M, Pokrovskii M, Ding Y, Yi R, Au C, Harrison OJ, Galan C, Belkaid Y, Bonneau R, Littman DR. c-MAF-dependent regulatory T cells mediate immunological tolerance to a gut pathobiont. Nature. 2018;554:373–7.

34. Mai V, McCrary QM, Sinha R, Glei M. Associations between dietary habits and body mass index with gut microbiota composition and fecal water genotoxicity: an observational study in African American and Caucasian American volunteers. Nutr J. 2009;8:49.

35. Mann E, Schmitz-Esser S, Zebeli Q, Wagner M, Ritzmann M, Metzler-Zebeli BU. Mucosa-associated bacterial microbiome of the gastrointestinal tract of weaned pigs and dynamics linked to dietary calcium-phosphorus. PLoS One. 2014;9: e86950.

36. Gomes JM, Costa JA, Alfenas RC. Could the beneficial effects of dietary calcium on obesity and diabetes control be mediated by changes in intestinal microbiota and integrity? Br J Nutr. 2015;14:1756–65.

37. Zhu Y, Michelle LT, Jobin C, Young HA. Gut microbiota and probiotics in colon tumorigenesis. Cancer Lett. 2011;309:119–27.

38. Venugopal PG, Nutman TB, Semnani RT. Activation and regulation of toll-like receptors (TLRs) by helminth parasites. Immunol Res. 2009;43:252–63.

39. Broadhurst MJ, Ardeshir A, Kanwar B, Mirpuri J, Gundra UM, Leung JM, Wiens KE, Vujkovic-Cvijin I, Kim CC, Yarovinsky F, Lerche NW, McCune JM, Loke P. Therapeutic helminth infection of macaques with idiopathic chronic diarrhea alters the inflammatory signature and mucosal microbiota of the colon. PLoS Pathog. 2012;8:e1003000.

40. Scotland RS, Stables MJ, Madalli S, Watson P, Gilroy DW. Sex differences in resident immune cell phenotype underlie more efficient acute inflammatory responses in female mice. Blood. 2011;118:5918–27.

41. Klein SL, Flanagan KL. Sex differences in immune responses. Nat Rev Immunol. 2016;16:626–38.

42. Scharsack JP, Koch K, Hammerschmidt K. Who is in control of the stickleback immune system: interactions between *Schistocephalus solidus* and its specific vertebrate host. Proc Biol Sci. 2007;274:3151–8.

43. Barber I, Scharsack JP. The three-spined stickleback-*Schistocephalus solidus* system: an experimental model for investigating host-parasite interactions in fish. Parasitology. 2010;137: 411–24.

44. Hendry AP, Peichel CL, Matthews B, Boughman JW, Nosil P. Stickleback research: the now and the next. Evol Ecol Res. 2013;15:111–41.

45. Houlden A, Hayes KS, Bancroft AJ, Worthington JJ, Wang P, Grencis RK, Roberts IS. Chronic *Trichuris muris* Infection in C57BL/6 mice causes significant changes in host microbiota and metabolome: Effects reversed by pathogen clearance. PLoS One. 2015;10:e0125945.

46. Whary MT, Muthupalani S, Ge Z, Feng Y, Lofgren J, Shi HN, Taylor NS, Correa P, Versalovic J, Wang TC, Fox JG. Helminth co-infection in *Helicobacter pylori* infected INS-GAS mice attenuates gastric premalignant lesions of epithelial dysplasia and glandular atrophy and preserves colonization resistance of the stomach to lower bowel microbiota. Microbes Infect. 2014;16:345–55.

47. Khan AR, Fallon PG. Helminth therapies: translating the unknown unknowns to known knowns. Int J Parasitol. 2013;43:293–9.

48. Whitacre, C.C. Sex differences in autoimmune disease. Nat Immunol. 2011;9:777–80.

49. Chiaroni-Clarke RC, Munro JE, Ellis JA. Sex bias in paediatric autoimmune disease - Not just about sex hormones? J Autoimmun. 2016;69:12–23.

50. Weber JN, Steinel NC, Shim KC, Bolnick DI. Recent evolution of extreme cestode growth suppression by a vertebrate host. Proc Natl Acad Sci USA. 2017;114:6575–80.

51. Lohman BK, Steinel NC, Weber JN, Bolnick DI. Gene expression contributes to the recent evolution of host resistance in a model host parasite system. Front Immunol. 2017;8:1071.

52. Smith CC, Snowberg LK, Gregory CJ, Knight R, Bolnick DI. Dietary input of microbes and host genetic variation shape among-population differences in stickleback gut microbiota. ISME J. 2015;9:2515–26.

53. Schloss PD, Westcott SL, Ryabin T, Hall JR, Hartmann M, Hollister EB, Lesniewski RA, Oakley BB, Parks DH, Robinson CJ, Sahl JW, Stres B, Thallinger GG, Van Horn DJ, Weber CF. Introducing mothur: open-source, platform-independent, community-supported software for describing and comparing microbial communities. Appl Environ Microbiol. 2009;5:7537–41.

54. Kozich JJ, Westcott SL, Baxter NT, Highlander SK, Schloss PD. Development of a dual-index sequencing strategy and curation pipeline for analyzing amplicon sequence data on the MiSeq Illumina sequencing platform. Appl Environ Microbiol. 2013;79:5112–20.

55. Quast C, Pruesse E, Yilmaz P, Gerken J, Schweer T, Yarza P, Peplies J, Glöckner FO. The SILVA ribosomal RNA gene database project: improved data processing and web-based tools. Nucleic Acids Res. 2013;41:D590–6.

56. Cole JR, Wang Q, Fish JA, Chai B, McGarrell DM, Sun Y, Brown CT, Porras-Alfaro A, Kuske CR, Tiedje JM. Ribosomal Database Project: data and tools for high throughput rRNA analysis. Nucleic Acids Res. 2014;42:D633–42.

57. McMurdie PJ, Holmes S. Phyloseq: an R package for reproducible interactive analysis and graphics of microbiome census data. PLoS One. 2013;8:e61217.

58. Peterson BK, Weber JN, Kay EH, Fisher HS, Hoekstra HE. Double digest RADseq: an inexpensive method for de novo SNP discovery and genotyping in model and non-model species. PLoS One. 2012;7:e37135.

59. Stuart YE, Veen T, Weber JN, Hanson D, Ravinet M, Lohman BK, Thompson CJ, Tasneem T, Doggett A, Izen R, Ahmed N, Barrett RDH, Hendry AP, Peichel CL, Bolnick DI. Contrasting effects of environment and genetics generate a continuum of parallel evolution. Nat Ecol Evol. 2017;1:158.

60. Broman KW, Wu H, Sen S, Churchill GA. R/qtl: QTL mapping in experimental crosses. Bioinformatics. 2003;19:889–90.

