## Supplemental Figs and Tables for "Host sex and genotype modify the gut microbiome response to helminth infection"

This file includes: Figures S1 to S9 and Tables S1 to S5.

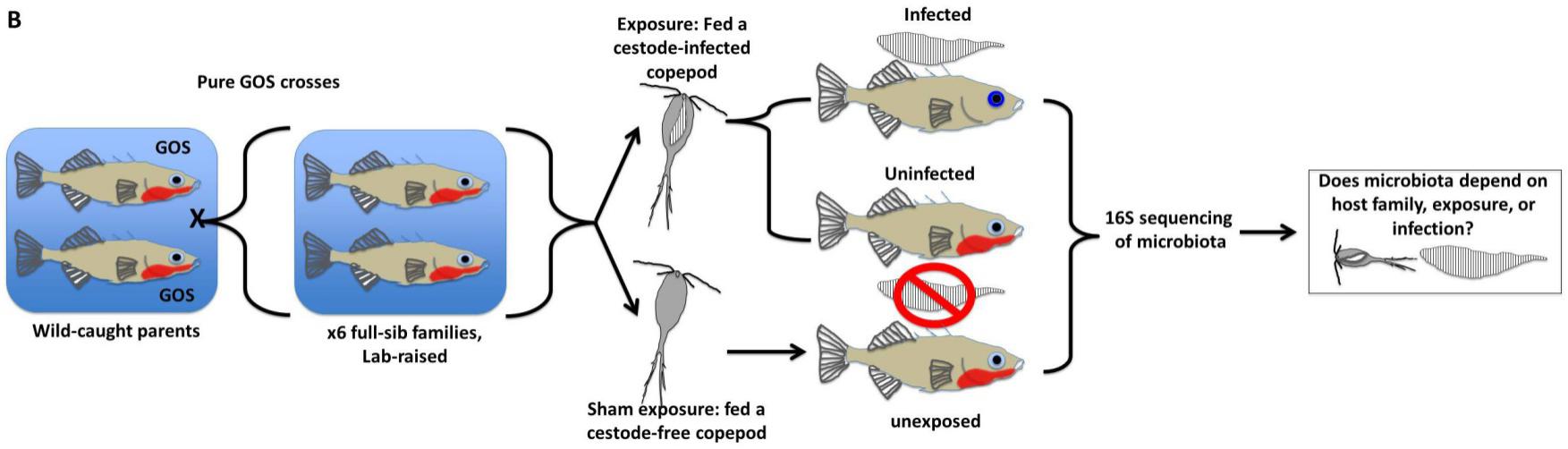

**a**

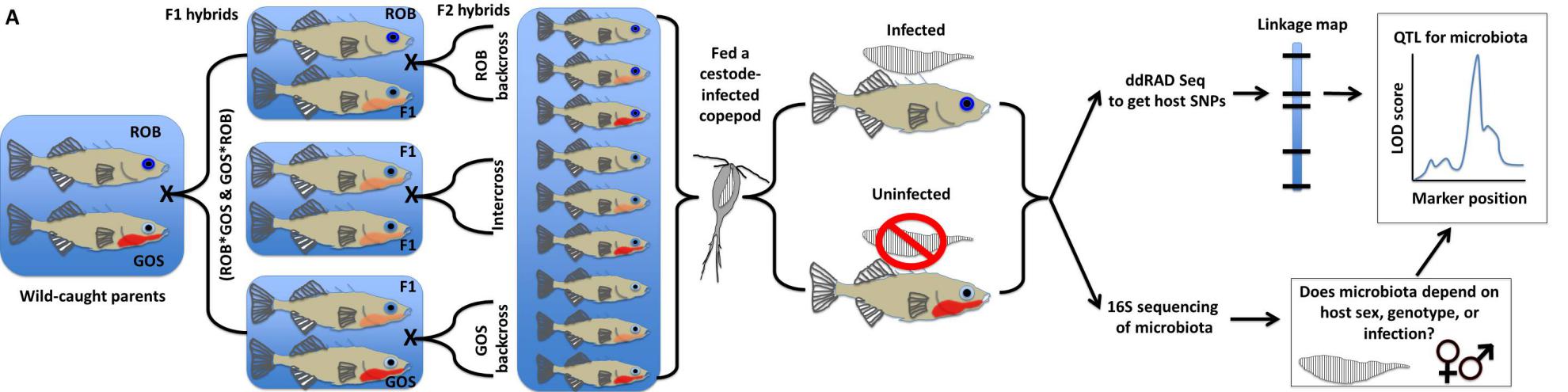

**b**

**Figure S1** Diagrams of study design**.** (a) Adult GOS (Gosling Lake) pure-bred fish drawn from six full-sibling families were fed with copepods with/without *S.solidus*. Three categories of fish within each family: unexposed controls, exposed-but-uninfected controls, and infected fish were obtained and used for subsequent microbiome analysis. (b) The sticklebacks derived from F2 hybrid crosses (intercrosses and both reciprocal backcrosses) between two lake populations (ROB(Roberts Lake) and GOS) of stickleback were exposed to *S.solidus*. A total of 693 fish were used for subsequent microbiome and QTL analysis. b)

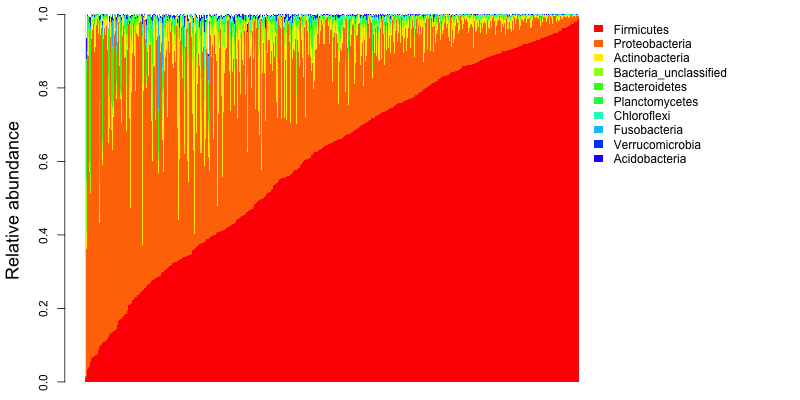

**Figure S2** Phylum relative abundance varies dramatically among individual stickleback. Only the 10 most common Phylum classifications are plotted here. Fish (x axis) are sorted in order of increasing relative abundance of Firmicutes.

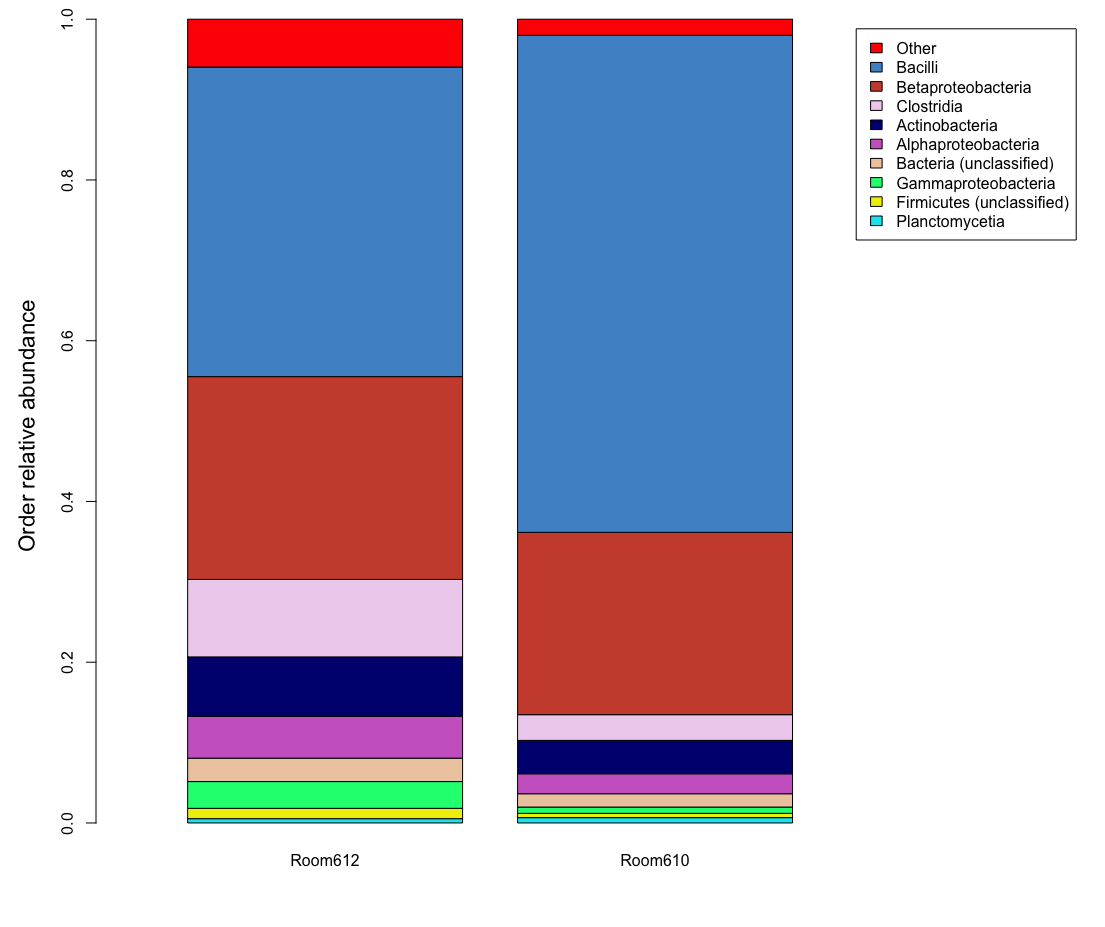

**Figure S3** The gut microbiota composition of the fish reared in Room 612 and 610.  Taxa summary plots at Order level, and data represents mean relative abundance.

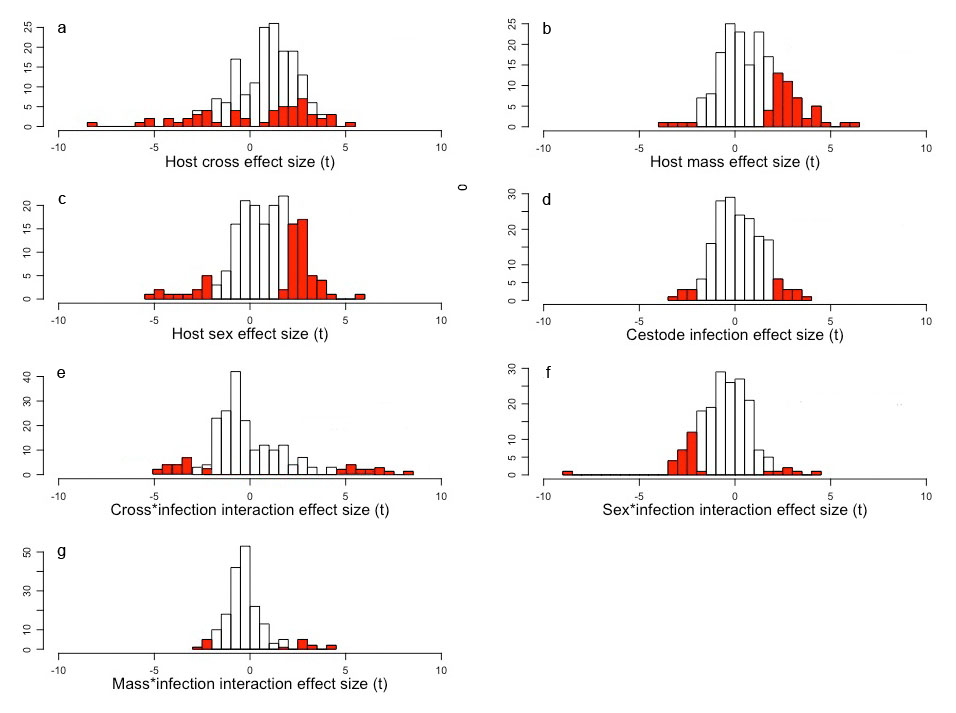

**Figure S4** Histograms of effect sizes of host (fish) cross type, host mass, host sex, infection status, and interactions between infection and cross or sex using quasibinomial GLMs with t statistics of regression coefficients as estimates. Here we plot red shaded areas denote statistically significant effects at a cutoff of *P*<0.05. Note that there is a strong excess of significant results at this threshold above the null expectation of a 5% false positive rate (all χ2 tests *P*<0.0001). (a) The relative abundance of 29.5% of Families depended on host cross type (*P*<0.05).(b) The relative abundance of 27.1% of Families depended on host host mass (*P*<0.05).(c) The relative abundance of 32.5% of Families depended on host host sex (*P*<0.05).(d) The relative abundance of 11.0% of Families depended on infection status (*P*<0.05).(e) The relative abundance of 19.9% of Families depended on host cross*infection interaction (*P*<0.05).(f) The relative abundance of 17.1% of Families depended on host sex*infection interaction (*P*<0.05).(g) The relative abundance of 8.5% of Families depended on host mass*infection interaction (*P*<0.05). Generalized linear model results for all common Families (found in at least 20 fish) are summarized in File S2.

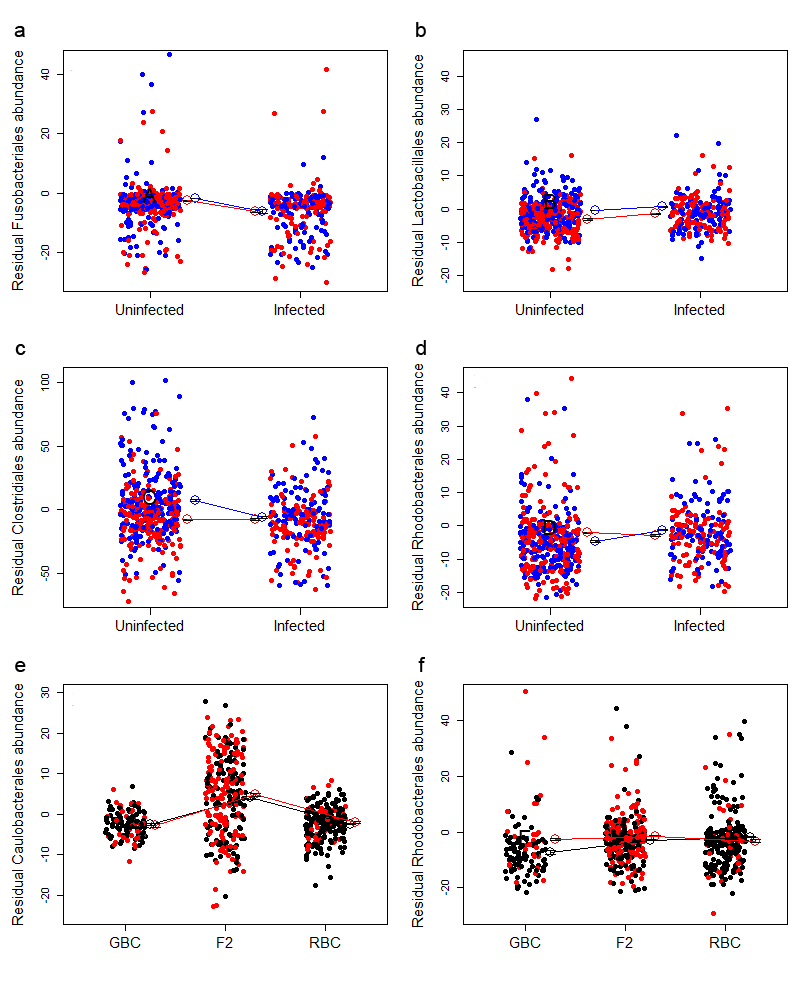

**Figure S5** Examples of how the gut microbiome composition depends on host sex, genotype (cross), and infection status, focusing on the relative abundance of microbial. Orders with all data points after statistically removing noise due to differences between rearing rooms. The same broad trends as Fig. 3 were observed, but this plot showed the small effect size relative to high among-individual variation. (a) Fusobacteriales, (males =blue and females =red). (b) Lactobacillales. (c) Clostridiales. (d) Rhodobacterales. (e) Caulobacterales (black = uninfected, red = infected). (f) Rhodobacterales.

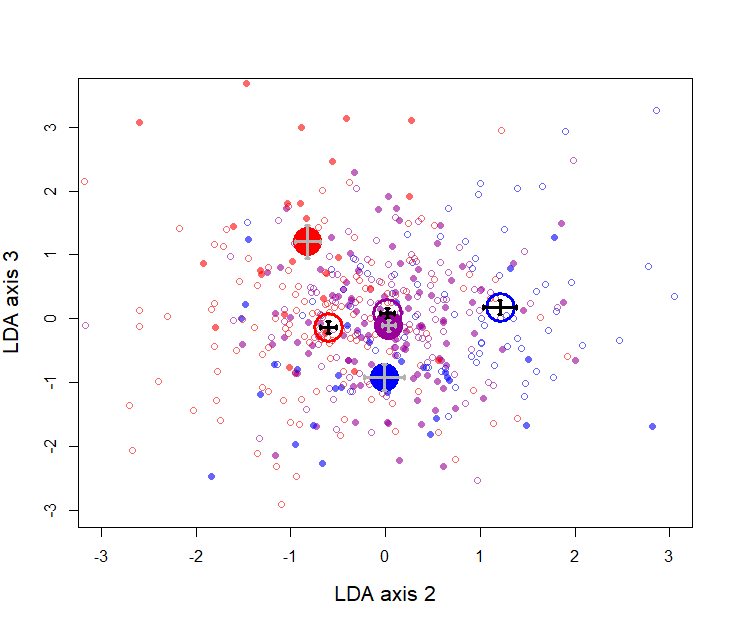

**Figure S6** Linear discriminant analysis of the top 50 unweighted PCoA axes of microbial composition reveals effects of host cross type and infection status. LDA2 (16% of variation) separates Roberts from Gosling backcross fish, with F2 hybrids being intermediate (blue = Gosling backcross, purple = F2, red = Roberts backcross) and infection status (open/filled circles). LDA3 explains 7% of the microbial variation and is most strongly associated with infection status, but in a manner that depends on host cross: the three host crosses are on average almost identical along LDA3 when uninfected (open circles), but diverge when infected (filled cycles) with F2s intermediate as expected from additive genetic control. Raw points are shown in faded colors, overlain by larger darker circles representing bivariate averages with ±1 s.e. error bars.

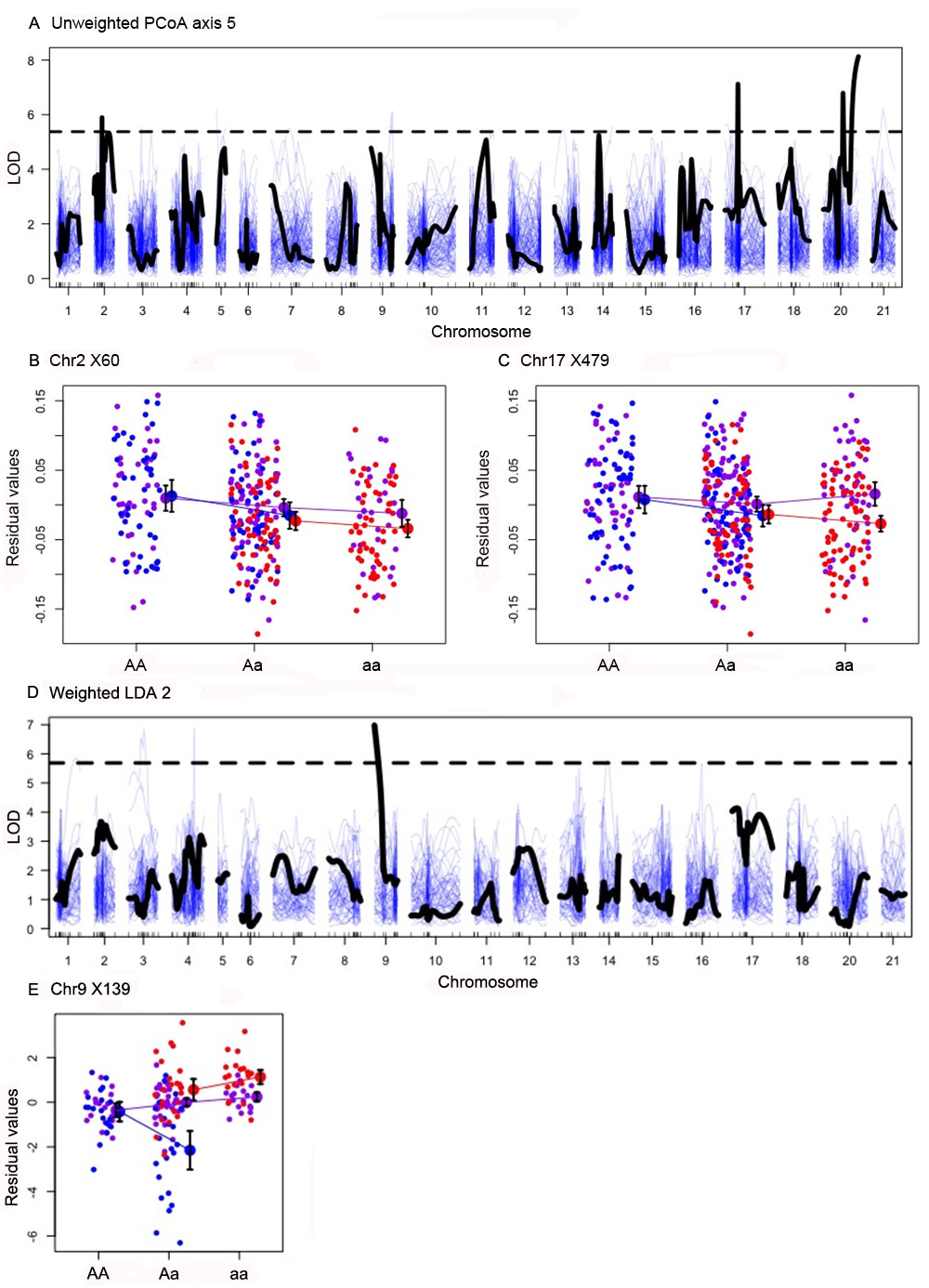
**Figure S7** Results of QTL mapping of microbial variation with unweighted PCoA axis 5/weighted PCoA axes 1-50 based on cross type. Analyses were run separately for each cross, with rearing room as an additive covariate, then the LOD scores from the three maps were summed. The observed summed LOD scores are plotted in black for each linkage group, measuring statistical association between the focal trait and the chromosomal region. Marker locations are indicated as tick marks along the horizontal axis. Thin blue lines represent null summed LOD scores from within-cross permutations of traits. The horizontal dashed line indicates the upper 99.99% quantile for the null LOD scores. (a-c) We focus here on QTL mapping of the resulting unweighted PCoA axis 5. Three QTL exceed this threshold, on Chr2, Chr 17, and Chr 20. Two of these are plotted in the lower panels (left, locus X60 on Chr2, genotype *P* = 0.00016; right, locus X479 on Chr17, genotype *P*=0.0386). The y axis in these effect plots are the residuals from a regression of unweighted PCoA5 on rearing room. We plot the raw data (color coded by cross, blue = Gosling backcross, purple = F2, red = Roberts backcross). For each genotype within each cross we also plot the means and one standard error bars (offset slightly to the right). (d-e) We focus here on QTL mapping of the resulting LDA axis 2 which exhibits additive effects of host genotype (e.g., F2 hybrids are between the backcrosses). No significant QTL was found for LDA axis 1, in which F2 hybrids are extreme relative to the two backcrosses implying non-additive genetic control which our QTL analysis is not well suited to detect. A single QTL is observed on Chr9 that exceeds the 99.99% threshold of null LOD values. In a linear model with rearing room as a covariate, the genotype at this locus is significantly associated with this microbial discriminant axis (*P*=0.002).

c

a

b

d

e

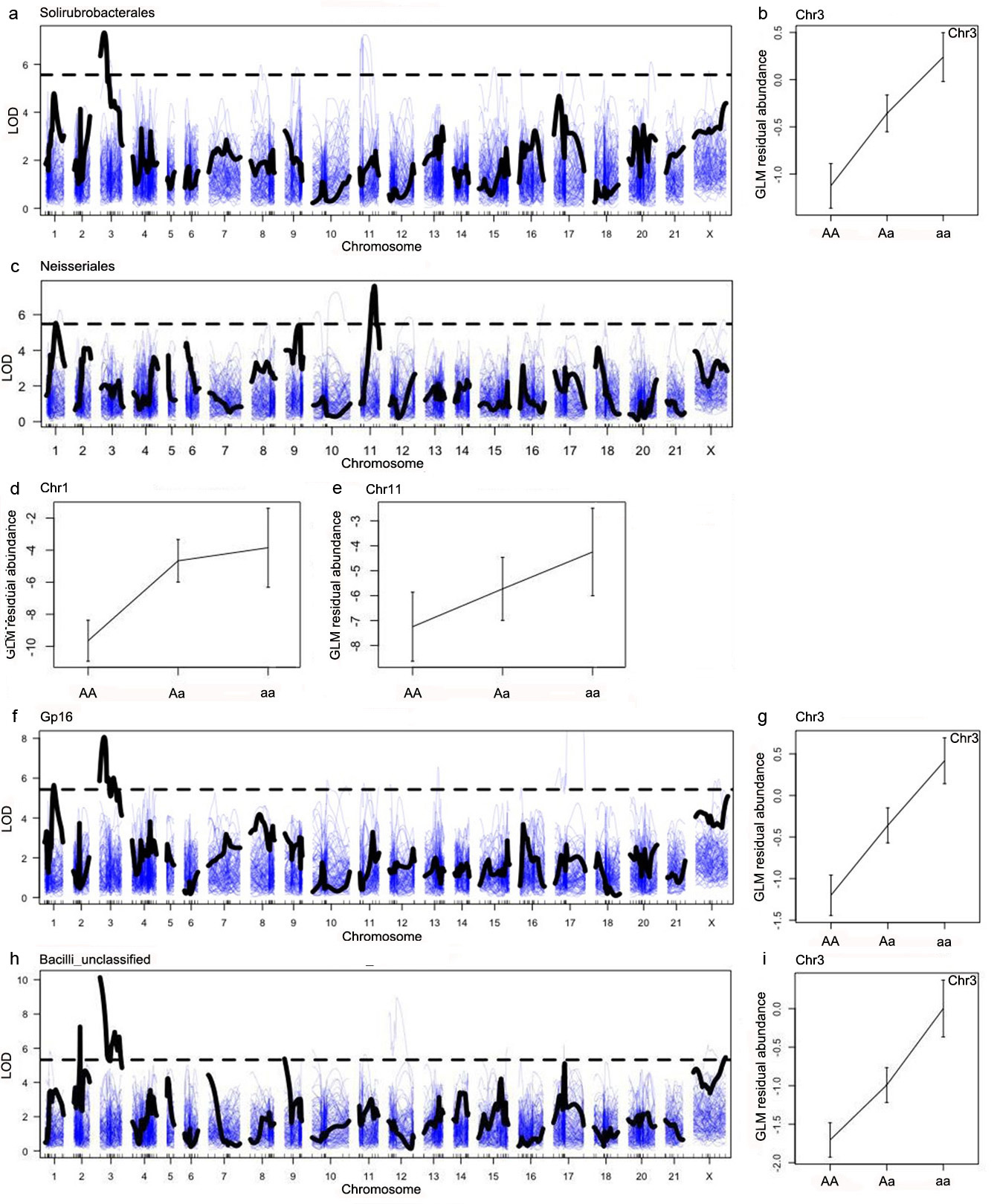

**Figure S8** Results of QTL mapping of the relative abundance of four microbial Orders. (a-b) QTL mapping of the relative abundance of Solirubrobacterales. An effect plot is provided for the single strong QTL on Chr3 (*P*=0.006). (c-e) QTL mapping of the relative abundance of Neisseriales. An effect plot is provided for the QTLs on Chr1 and Chr11 (*P*=0.070, 0.0092 respectively). (f-g) QTL mapping of the relative abundance of Gp16. An effect plot is provided for the QTL on Chr3 (*P*=0. 0.0054). The QTL on Chr1 does not quite approach significance (*P*=0.183) in a separate linear model with rearing room as a covariate. (h-i) QTL mapping of the relative abundance of Bacilli_unclassified. An effect plot is provided for the QTL on Chr3 (*P*=0.0043).

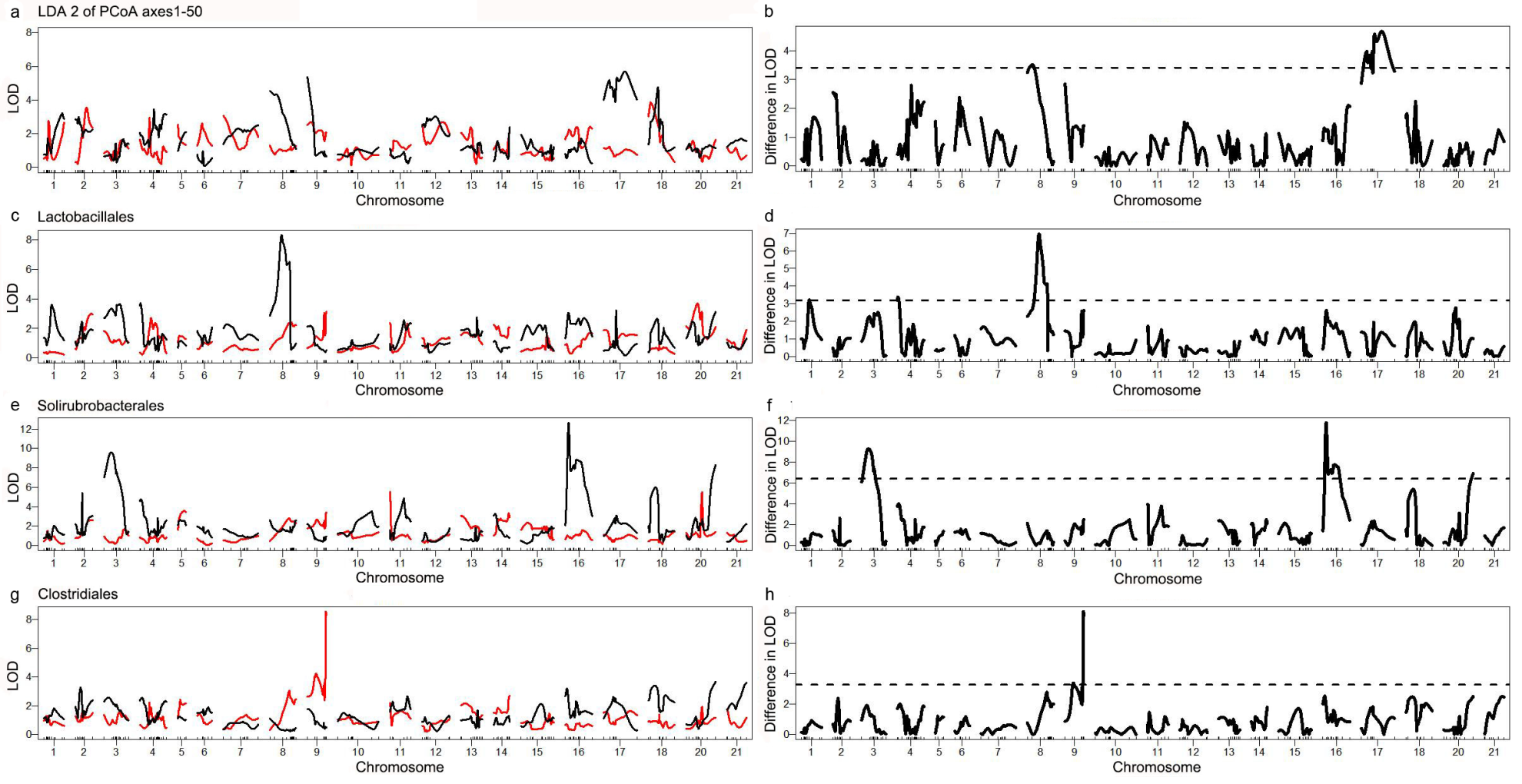

**Figure S9** Infection-dependent QTL for overall microbial community structure as measured by the second linear discriminant axis (LDA2) of PCoA axes 1-50 and the relative abundance of microbial Orders. In the left panel we plot LOD scores for infected (red) and uninfected (black) stickleback using rank-based mapping to obtain LOD values. The right panel plots the absolute magnitude difference in LOD scores between infected versus uninfected fish, as a test of support for a QTL genotype by infection interaction. To obtain confidence intervals for this LOD difference, we randomized fish within each cross type and infection status. We then re-ran QTL analyses within each cross and infection combination. For each infection group we summed the LOD scores for ROB backcross, F2, and GOS backcross fish. Then to obtain a null LOD difference, we took the absolute value of the difference between the LOD for all infected fish minus the LOD for all uninfected fish. We repeated this null calculation 1000 times (thin blue lines). We considered a QTL-infection interaction to be significant if the observed LOD difference (thick black line) exceeded the 99.99% quantile for the null LOD differences (dashed horizontal line). (a-b) For overall microbial community structure (LDA2 PCoA axes 1-50), we observe strong QTL for microbial composition on Chr8 and Chr17, but only in uninfected fish. The difference in LOD scores between infected versus uninfected fish is highly significant for Chr17, and marginally so on Chr8. (c-d) For the relative abundance of Lactobacillales, we observe a strong QTL for Lactobacillales abundance on Chr8, but only for uninfected fish. There is no comparable QTL for infected fish (red). The difference between their respective LOD scores on Chr8 substantially exceeds the 99.99% quantile for null values. We conclude that there is a genetic variant on Chr8 that regulates Lactobacillales abundance in the absence of *S.solidus* infection, but that helminth infection eliminates that effect. (e-f) For Solirubrobacterales relative abundance, we observe strong QTL for microbial abundance on Chr3 and Chr16. But, both QTL are only observed in uninfected fish (black lines) and there is no significant QTL for this microbial Order in infected fish. The difference in LOD scores for Chr3 and Chr16 are highly significant when compared with null expectations from a permutation analysis. (g-h) For Clostridiales relative abundance, we observe strong QTL for microbial abundance on Chr9, but only in infected stickleback. The difference in LOD scores for Chr9 is highly significant when compared with null expectations from a permutation analysis. We conclude that *S.solidus* infection facilitates expression of a genetic variant on Chr9 that can alter Clostridiales abundance, but which is normally inactive without the tapeworm.

**Table S1.** Sample sizes of stickleback used in a separate experiment in which we experimentally exposed lab-raised fish from one lake (Gosling Lake) to *S.solidus* population.

|  | Gosling Lake stickleback family | | | | |  |
| --- | --- | --- | --- | --- | --- | --- |
|  | GG8 | GG12 | GG13 | GG15 | GG16 | GG17 |
| Control | 2 | 2 | 2 | 1 | 1 | 2 |
| Exposed but uninfected | 1 | 2 | 2 | 2 | 1 | 2 |
| Infected | 1 | 3 | 1 | 1 | 3 | 1 |

**Table S2**. *Adonis* analysis of weighted and unweighted UniFrac distances between infected, uninfected, and control stickleback from a separate experiment in which we experimentally exposed lab-raised fish from one lake (Gosling Lake) to *S.solidus* population, as well as alpha diversity.

|  | Dependent variable | Df | Sum of Squares | F | r^2^ | P |
| --- | --- | --- | --- | --- | --- | --- |
| Model 1 | Unweighted UniFrac | | |  |  |  |
| Infection |  | 1 | 0.1216 | 0.5067 | 0.0157 | 0.9914 |
| Family |  | 5 | 2.3515 | 1.9605 | 0.3031 | <0.0001 |
| Infection*Family |  | 5 | 0.9673 | 0.8065 | 0.1247 | 0.9331 |
| Residuals |  | 18 | 4.3179 |  | 0.5566 |  |
| Model 2 | Weighted UniFrac | |  |  |  |  |
| Infection |  | 1 | 0.0232 | 0.2983 | 0.0088 | 0.9708 |
| Family |  | 5 | 0.9052 | 2.3250 | 0.3442 | 0.0033 |
| Infection*Family |  | 5 | 0.3001 | 0.7709 | 0.1141 | 0.7827 |
| Residuals |  | 18 | 1.4016 |  | 0.5329 |  |
| Model 3 | Unweighted UniFrac | | |  |  |  |
| Exposure |  | 1 | 0.2421 | 1.1503 | 0.0312 | 0.2589 |
| Family |  | 5 | 2.3623 | 2.2444 | 0.3045 | <0.0001 |
| Exposure*Family |  | 5 | 1.3646 | 1.2965 | 0.1759 | 0.0363 |
| Residuals |  | 18 | 3.7892 |  | 0.4884 |  |
| Model 4 | Weighted UniFrac | |  |  |  |  |
| Exposure |  | 1 | 0.0346 | 0.5254 | 0.0131 | 0.7998 |
| Family |  | 5 | 0.8984 | 2.7315 | 0.3416 | 0.0007 |
| Exposure*Family |  | 5 | 0.5131 | 1.5602 | 0.1951 | 0.0631 |
| Residuals |  | 18 | 1.1841 |  | 0.4502 |  |

**Table S3**. Sample sizes of stickleback used in this study for sequencing (*N*=693). In parentheses we provide the number of stickleback experimentally exposed to *S.solidus* (*N*=711).

| Room | Exposure outcome | Stickleback cross type | | |
| --- | --- | --- | --- | --- |
|  |  | Gosling backcross | F2 intercross | Roberts backcross |
| Room 612 | Cestode failed | 0 | 68 (69) | 37 (37) |
| Room 612 | Cestode established | 0 | 115 (115) | 17 (17) |
| Room 610 | Cestode failed | 107 (109) | 70 (71) | 155 (157) |
| Room 610 | Cestode established | 42 (52) | 37 (37) | 45 (47) |

**Table S4**. Gut microbiome sample sizes after rarefaction to 500/2000/4000 sequences each, removing individuals (43/165/250) with insufficient coverage.

| Room | Exposure outcome | Stickleback cross type | | |
| --- | --- | --- | --- | --- |
|  |  | Gosling backcross | F2 intercross | Roberts backcross |
| Room 612 | Cestode failed | 0 | 67/55/47 | 34/19/13 |
| Room 612 | Cestode established | 0 | 110/89/77 | 15/12/9 |
| Room 610 | Cestode failed | 103/95/73 | 67/50/40 | 137/114/98 |
| Room 610 | Cestode established | 39/37/34 | 35/26/22 | 43/31/30 |

**Table S5**. Results of a MANOVA testing effects of infection status, host sex, host mass, host cross (Gosling backcross, F2, Roberts backcross) and interactions among these variables on microbial community composition (weighted or unweighted PCoA axes 1-50).

| Model effect |  | Unweighted PCoA 1-50 | | |  | Weighted PCoA 1-50 | | |
| --- | --- | --- | --- | --- | --- | --- | --- | --- |
|  | Effect Df | Pillai's trace | F | F Df | *P* | Pillai's trace | F | *P* |
| Host cross | 2 | 0.986 | 8.865 | 100 ; 912 | 0.0000 | 0.750 | 5.471 | 0.0000 |
| Infection | 1 | 0.183 | 2.043 | 50 ; 455 | 0.0001 | 0.134 | 1.407 | 0.0403 |
| Host sex | 1 | 0.202 | 2.303 | 50 ; 455 | 0.0000 | 0.200 | 2.278 | 0.0000 |
| Host mass | 1 | 0.396 | 5.959 | 50 ; 455 | 0.0000 | 0.293 | 3.771 | 0.0000 |
| Host cross * infection | 2 | 0.291 | 1.551 | 100 ; 912 | 0.0008 | 0.243 | 1.264 | 0.0483 |
| Host cross * sex | 2 | 0.212 | 1.080 | 100 ; 912 | 0.2872 | 0.227 | 1.165 | 0.1388 |
| Host cross * mass | 2 | 0.443 | 2.598 | 100 ; 912 | 0.0000 | 0.346 | 1.911 | 0.0000 |
| Infection * sex | 1 | 0.124 | 1.287 | 50 ; 455 | 0.0983 | 0.099 | 0.995 | 0.4867 |
| Infection * mass | 1 | 0.129 | 1.351 | 50 ; 455 | 0.0616 | 0.108 | 1.105 | 0.2966 |
| Sex * mass | 1 | 0.101 | 1.021 | 50 ; 455 | 0.4391 | 0.128 | 1.336 | 0.0692 |
| Residuals | 504 |  |  |  |  |  |  |  |
